# Chromosomal inversions mediated by tandem insertions of transposable elements

**DOI:** 10.1101/2025.01.09.631634

**Authors:** Robin Aasegg Araya, William B Reinar, Ole K Tørresen, Clément Goubert, Tara J Daughton, Siv Nam Khang Hoff, Helle T Baalsrud, Marine Servane Ono Brieuc, Anna Komisarczuk, Sissel Jentoft, José Cerca, Kjetill S Jakobsen

## Abstract

Chromosomal inversions play a crucial role in evolution by influencing phenotypes through the linkage of co-adapted alleles. While inversions have been found across a large number of taxa, mapping and characterizing inversion breakpoint regions remains challenging, often due to the presence of complex tandem repeats and transposable elements (TEs). Here, we identify and quantify TEs in the breakpoints of the four large-scale inversions previously reported in Atlantic cod, leveraging on three high-quality long-read-based reference genome assemblies for the Norwegian Coastal cod, the Northeast Arctic cod and Celtic cod ecotypes. We detected a significant enrichment of TE orders and superfamilies with terminal inverted repeats (TIRs) within the inversion breakpoints of chromosomes 1, 7 and 12. Notably, we discovered a tandem accumulation of miniature inverted-repeat transposable elements (MITEs) belonging to a family of *hAT* transposons, exclusively residing in the breakpoints of the inverted haplotype on chromosomes 1 and 7 found in the Northeast Arctic cod. The accumulation of tandemly arranged TEs in breakpoint regions suggests that they have driven the appearance of inversions through ectopic recombination, further supporting the potential of TEs in facilitating chromosomal reorganizations with large evolutionary implications.

## Introduction

Genetic variation is essential for evolution and can arise from various sources, including single-point mutations, transposable element (TE) activity, and structural variants like chromosomal inversions (Sætre & Ravinet, 2019). In particular, inversions have been shown to include sets of co-adapted allelic variants within the regions they encompass (Wellenreuther & Bernatchez, 2018; Mérot et al., 2020; Gutiérrez-Valencia et al., 2021; Villoutreix et al., 2021), that can potentially have an important effect on the fitness of the individuals. As a result, inversions have important evolutionary implications for the processes of adaptation (Faria et al., 2019; Mérot et al., 2020) and speciation (Wellenreuther & Bernatchez, 2018). Examples of the impact of inversions include variation in mimicry patterns associated with local adaptation in the butterfly species *Heliconius numata* (Jay et al., 2018), different life history strategies related to mating in ruffs (*Calidris pugnax*) (Lamichhaney et al., 2016), and migratory spawning behavior in Atlantic cod (*Gadus morhua*) (Berg et al., 2016).

Despite the growing appreciation of the evolutionary impact of chromosomal inversions, the molecular mechanisms by which inversions originate are yet to be fully explored. Double-stranded breaks on the same chromosome repaired by non-homologous end joining (NHEJ) can result in an inversion (Ranz et al., 2007; Guillén & Ruiz, 2012). Inversions can also result from ectopic recombination (Ling & Cordaux, 2010; Guillén & Ruiz, 2012), in which recombination happens between similar sequences that are present in different genomic regions, such as homologous TEs in opposite directions on a chromosome. Identifying the underlying mechanisms responsible for the generation of inversions, however, can be challenging (Faria et al., 2019), and the relative contributions of NHEJ and ectopic recombination have not been investigated in detail, with few exceptions (Ranz et al., 2007; Guillén & Ruiz, 2012). The advent of high-quality chromosome-level genomes (Peona et al., 2018; The Darwin Tree of Life Project Consortium et al., 2022) and new bioinformatic tools provide promising avenues for understanding the mechanisms that generate inversions.

The Atlantic cod (*Gadus morhua*) genome enharbour four large chromosomal inversions that have been linked to adaptations to environmental and ecological conditions such as temperature, light, oxygen, salinity at different geographical sites (Berg et al., 2015, 2017; Barth et al., 2017, 2019), and distinguish the two ecotypes: the migratory North East Arctic cod (NEAC) and the stationary Norwegian Coastal Cod (NCC) (Berg et al., 2016; Matschiner et al., 2022). In a recent study, Matschiner et al. (2022) demonstrated that the migratory ecotype (NEAC) is associated with the derived (inverted) haplotype of the inversions on chromosomes 1 and 7, while the ancestral (non-inverted) haplotype is predominant in the non-migratory ecotype (NCC) (Matschiner et al., 2022). Conversely, the non-migratory NCC is associated with the derived (inverted) haplotypes on chromosomes 2 and 12, whereas the ancestral (non-inverted) arrangements are mostly found in the migratory ecotype (Matschiner et al., 2022). The four inversions originated at different ages from 0.40 to 1.66 million years ago (Matschiner et al., 2022), suggesting that they evolved within the Atlantic cod lineage.

The presence of these four large chromosomal inversions offers a unique opportunity to compare inversions and their evolution. Specifically, it is possible to pinpoint whether the different inversions have originated from similar molecular processes or not. One way to investigate this is to look at the inversion breakpoints. If NHEJ is involved, we expect to see no TE accumulation and the presence of short inverted duplications of non-repetitive DNA at the breakpoints. Conversely, if ectopic recombination is involved it is likely that an accumulation of similar sequences, such as homologous parts of related TEs, occurs in the breakpoint regions, as shown in previous studies (Cáceres et al., 1999; Zhang & Peterson, 2004; reviewed in Böhne et al., 2008; Delprat et al., 2009; Guillén & Ruiz, 2012; Sarilar et al., 2015; Sharma & Peterson, 2023). However, characterization of the breakpoint regions in Atlantic cod at the sequence level has not been done. Doing so will elucidate the underlying mechanisms and origins of inversions, which have often been overlooked (reviewed in Villoutreix et al., 2021).

Here, we perform a survey of TEs associated with the four chromosomal inversions on chromosomes 1, 2, 7, and 12 in Atlantic cod. Using synteny analyses and PacBio HiFi sequencing, we precisely locate inversion breakpoints and manually curate the TEs in breakpoint pairs for the NEAC and NCC ecotypes. We identify accumulations of specific TE families overlapping the breakpoints of all four inversions, with enrichments of certain TEs containing terminal inverted repeats (TIRs). We find tandemly arranged miniature inverted-repeat transposable elements (MITEs) related to the DNA transposon *hAT* superfamily in the breakpoints on chromosomes 1 and 7. Additional population-level analyses of the chromosome 2 inversion in 58 cod individuals reveal conserved breakpoint locations between individuals, also in terms of TE insertions. Overall, our findings suggest that TEs are directly involved in recombination events leading to inversion formation and thereby indirectly contribute to local adaptation.

## Results

### Inversion breakpoints are associated with accumulations of specific TE superfamilies

To determine the genomic coordinates of the inversions on chromosome 1, 2, 7 and 12, we used a combination of chromosome alignments and mapping of raw reads (see **Methods and Materials; Supplementary Table S1** and **Supplementary Figures S1-S4**) onto the NEAC, NCC and Celtic cod reference genome assemblies. This led us to identify each of the different breakpoint intervals to the scale of a few thousand base pairs (mean breakpoint interval size: 16,862 bp; SD: 21,417 bp; **Table 1**).

**Table 1.** Overview of breakpoint coordinates in the Atlantic cod inversions. All coordinates refer to the assembly carrying the derived (inverted) haplotype. The coordinates were determined by split read alignment and are provided as ranges.

| Chr | Breakpoint coordinates |  |  | Arrangement |  |
| --- | --- | --- | --- | --- | --- |
|  | A | B | C | Derived | Ancestral |
| 1<sup>a</sup> | 11,296,105-11,302,315 | 19,180,947-19,187,825 | 28,333,982-28,336,075 | NEAC | NCC |
| 2 | 21,848,272-21,858,616 | 26,151,340-26,163,504 | - | NCC | NEAC |
| 7 | 16,838,295-16,871,460 | 26,338,531-26,410,445 | - | NEAC | NCC |
| 12 | 638,394-646,752 | 13,659,852-13,660,483 | - | NCC | NEAC |

<sup>a</sup>The inversion on chromosome 1 is a double inversion with adjacent breakpoints (B).

Following identification of inversion breakpoints, we created a species-specific, manually curated TE library using an in-house pipeline (see **Materials and Methods; Supplementary Table S2** and **Supplementary Figure S5**). The resulting library masked 35.23% of the NEAC assembly, with 28.3% being interspersed repeats (**Supplementary Table S3**), and showed an even distribution of DNA elements, long terminal repeats (LTRs), and long interspersed nuclear elements (LINEs) with on average low divergence from their consensus sequences (mean divergence: 11.33%; *SD*: 11.65, see **Supplementary Figure S6**). Similar results were observed when masking the NCC and Celtic cod assemblies, indicating no substantial differences in TE annotations across the three cod ecotypes (**Supplementary Figure S6**). The TE annotations exhibited an NTE50 of 86,886 TEs (i.e., number of TE insertions required to annotate 50% of the genomic repeats) and LTE50 of 465 bp (i.e. the minimum length of annotated TEs needed to cover 50% of the genomic repeats), which is an improvement from the initial uncurated library, with shorter and more fragmented annotations (NTE50 = 123,454 TEs and LTE50 = 344 bp).

To investigate the relationship between TEs and inversion breakpoints, we used our TE library to assess TE enrichment around the breakpoints on chromosomes 1, 2, 7 and 12, comparing these regions to the overall chromosomal TE densities within sliding windows of 50 kb. We observed significantly elevated TE densities in the inversion breakpoints on chromosome 7 (NEAC) and chromosome 12 (NCC) compared to non-breakpoint regions on the same chromosomes (**Figure 1a** and **Supplementary Table S4**) (Student’s t-test with p-values: 5.42 x 10^-5^ and 6.4 x 10^-6^, respectively, excluding putative centromeric regions as estimated in **Supplementary Table S5** and **Supplementary Figures S7-S9**). In addition, the syntenic breakpoint-regions of the ancestral (non-inverted) haplotype on chromosome 12 in NEAC exhibited higher TE densities than the chromosomal average (p-value: 0.025). Chromosomes 1 and 2 did not display the same increase in general TE density around the breakpoints (**Figure 1a** and **Supplementary Figure S10a-d**).

**Figure 1:**
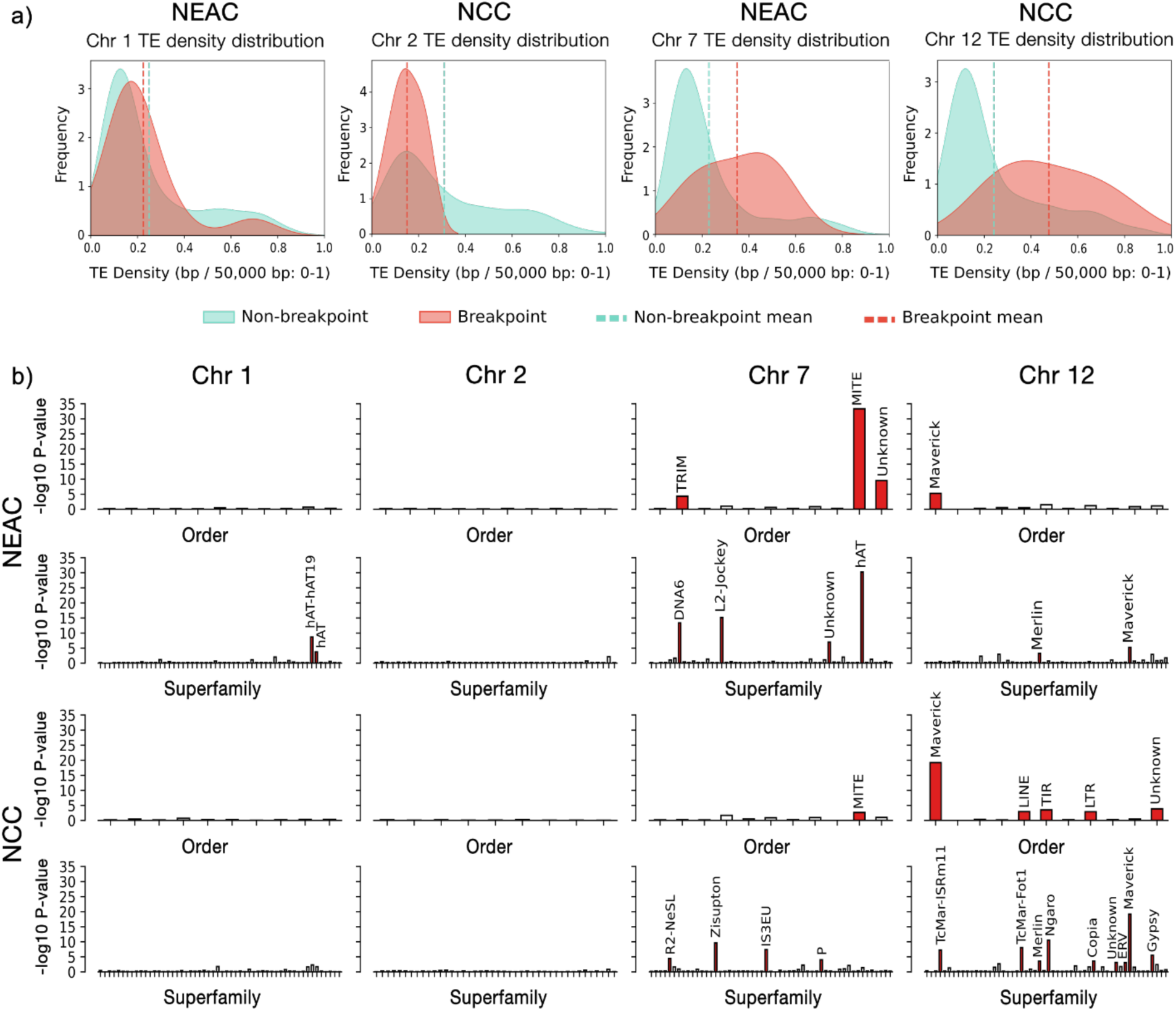
TE enrichment in breakpoints vs. non-breakpoint regions. **a,** Density distribution of TEs across breakpoint and non-breakpoint regions on chromosomes 1, 2, 7 and 12, using 50 kb non-overlapping sliding-windows. The red curve represents TE density within breakpoint regions, while density in non-breakpoint regions is shown in blue. Mean densities for breakpoint and non-breakpoint regions are indicated by red and blue dotted lines, respectively. **b,** TE orders and superfamilies associated with breakpoint regions using a Student’s t-test. TE orders and superfamilies are shown on the x-axis, and -log_10_ p-values on the y-axis for chromosomes 1, 2, 7 and 12 in NEAC and NCC. Red bars indicate TE orders and superfamilies with significantly elevated densities within breakpoint regions.

Further analysis of TE orders revealed above-average accumulations of specific TE orders within certain inversion breakpoints, where mean densities in the breakpoints exceeded 2 standard deviations above the chromosome-specific mean (excluding putative centromeric regions, **Supplementary Figures S10e-f**). In particular, we identified high densities of miniature inverted- repeat transposable elements (MITEs) in the breakpoints of the derived (inverted) haplotype on chromosome 7 in NEAC (breakpoint mean density: 6.5%; *SD*: 1.3% vs. chromosomal mean density: 0.4%; *SD*: 1.0%). Additionally, visual inspection of individual breakpoints showed an enrichment of class II DNA elements (density peaks above 2 *SD* of chromosomal mean) in the breakpoints of the derived (inverted) haplotypes on chromosomes 1 and 7 in NEAC, specifically of terminal inverted repeats (TIRs) and MITEs (**Supplementary Figure S10e**). We also observed elevated densities of retroelements with long terminal repeats (LTRs) within two of the breakpoints on chromosome 1 in NEAC. On chromosome 12, we found elevated levels of TIRs, retroelements, and Maverick elements in at least one of the breakpoints and syntenic breakpoint regions (NCC and NEAC, respectively) (**Supplementary Figure S10e-f**).

To better quantify the relationship between TE orders and inversion breakpoints, we performed statistical testing of TE order density differences between breakpoint and non-breakpoint regions within 50 kb sliding windows, using a Student’s t-test. This revealed a significant tendency for MITEs to accumulate in the breakpoints of the chromosome 7 inversion in NEAC (but not in the ancestral syntenic regions in NCC) (**Figure 1b**). Similarly, the breakpoints of the chromosome 12 inversion in NCC showed significantly elevated densities of Maverick elements, but to a lesser extent in the syntenic regions of NEAC (**Figure 1b**).

Finally, we extended this analysis to the TE superfamily level, performing the same density comparison within 50 kb sliding windows for breakpoints vs. non-breakpoint regions using a Student’s t-test. This yielded significant signals of enrichment for certain superfamilies of TIR elements (*hAT*-elements, DNA-6, and unspecified MITEs) in the breakpoints of the chromosome 7 inversion in NEAC, along with a superfamily of long interspersed nuclear elements (LINEs) (**Figure 1b** and **Supplementary Figure S10e**). For the chromosome 12 inversion in NCC, we again detected significantly elevated densities of an unspecified superfamily of Maverick elements within the breakpoints (**Figure 1b** and **Supplementary Figure S10e-f**).

### Closely related TEs occupy both breakpoints of all four inversions in Atlantic cod

Given the apparent association between inversion breakpoints and specific types of TEs, we wanted to investigate the sequence similarity of family-specific TE accumulations between breakpoint pairs (i.e., breakpoints of the same inversion). To capture the full TE content surrounding each breakpoint, we defined a breakpoint region of an inversion as the midpoint of each estimated breakpoint from **Table 1** and extended it by 50 kb in both directions to include flanking sequences, resulting in breakpoint regions of the same 100 kb size (see **Material and Methods**).

By dot-plotting each breakpoint pair, we identified multiple similar sequence fragments between breakpoints (sequence identity ≥ 25%) (**Supplementary Figure S11**). For all four derived (inverted) haplotypes, these fragments contained TE copies that were classified into the same family (**Figure 2**), explaining the sequence similarity between the breakpoints. On chromosome 1, we found three different TE families from the class II DNA elements that were present in the three breakpoints in NEAC—which carry the derived double inversion—whereas these TEs were absent from the ancestral syntenic regions in NCC (where we found short insertions of one family of DNA elements in two of the syntenic “breakpoints”) (**Figure 2**). A similar pattern was observed for the chromosome 12 inversion, where two families of class II DNA elements were found only in the derived (inverted) haplotype in NCC and not in the non-inverted haplotype in NEAC (**Figure 2**). On chromosome 2, both cod ecotypes showed a diversity of accumulated retroelements, primarily from the LTRs (**Figure 2**). Chromosome 7 exhibited insertions of various putative TE families (both LTRs and DNA elements) in the breakpoints of the derived (inverted) haplotype in NEAC, while two families of retroelements were present only in the syntenic breakpoint regions of NCC (**Figure 2**). Manual inspection of consensus sequences and insertions confirmed that the breakpoint insertions consistently represented the same or overlapping portions of their respective TE family consensus sequence (**Supplementary Table S6**).

**Figure 2:**
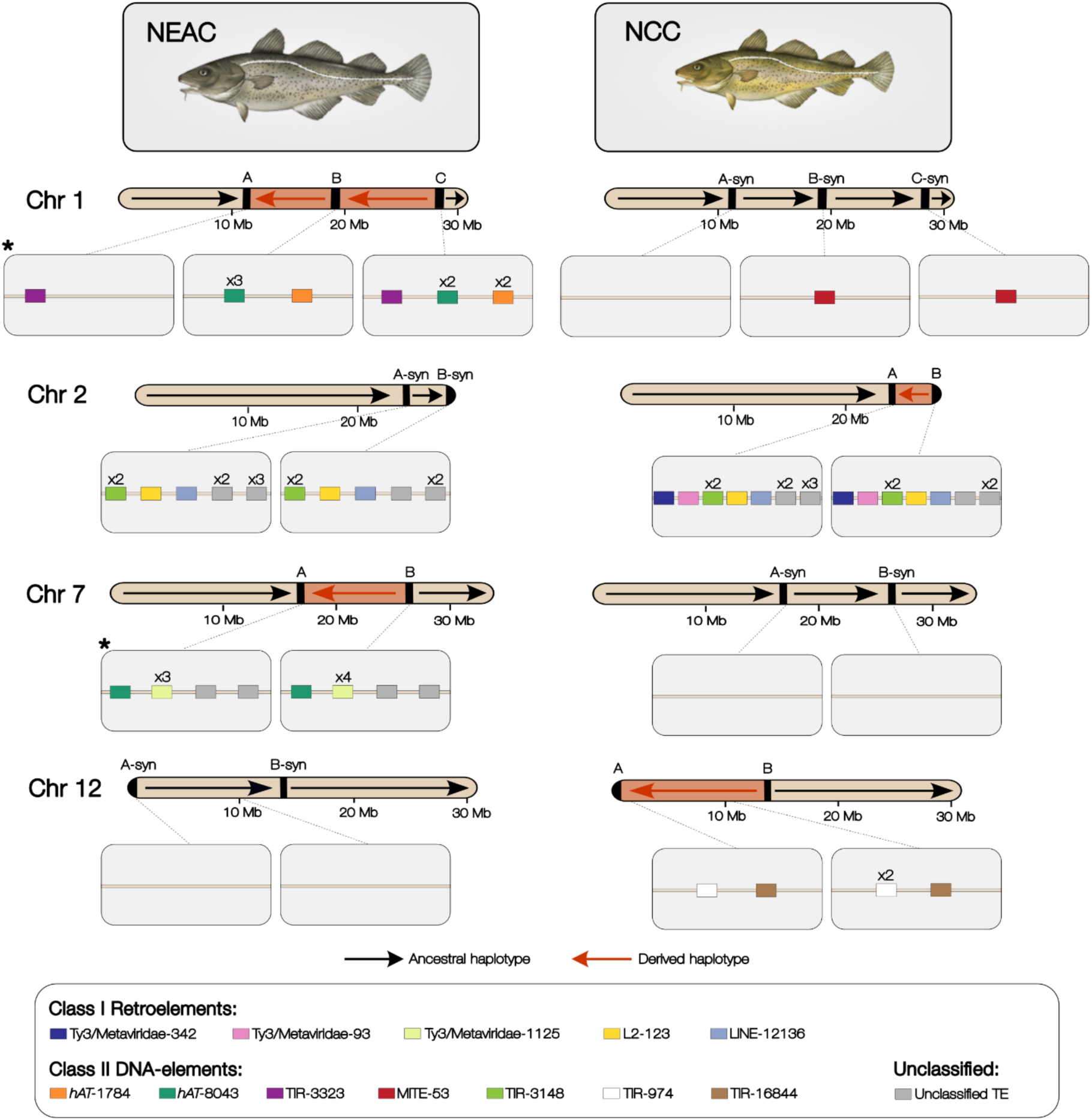
Transposable Elements (TEs) in the inversion breakpoints of Atlantic cod. TE families present in both breakpoints of an inversion in chromosomes 1, 2, 7 and 12 in NEAC and NCC are displayed as squares, with colors corresponding to TE families as indicated in the legend. Each chromosome shows breakpoint and syntenic breakpoint regions (midpoint ± 50 kb) within the light-gray boxes below. Black arrows on the chromosome cartoons show the ancestral (non-inverted) haplotype, red arrows show the derived (inverted) haplotype. Insertion lengths and orientations of the TE families are not shown. Multiple insertions are indicated with an x and number of copies. Chromosomes are drawn to scale, with sizes indicated in Mb. *Indicates breakpoints with a significantly higher occurrence of long TE insertions (>1,000 bp) of the same family, compared to randomly selected non-breakpoint regions (see **Supplementary Figure S13**).

The TE families that were present in breakpoint pairs (**Figure 2**) varied from 2 to 8 copies and exhibited diverse consensus lengths, ranging from approximately 284 bp to 9,034 bp (**Supplementary Table S6**). These TE families were not exclusively found in breakpoints (**Supplementary Figure S12**). Due to the accumulation of TEs from the same family in breakpoints, we compared the probability of finding such an accumulation of family-specific TEs in the breakpoint pairs *vs* random genomic regions. Specifically, we compared counts of observed family-specific TEs within both breakpoints of each inversion, to the average observed family-specific TEs occurring in 1,000 randomly selected region pairs on the same chromosome (**Supplementary Figure S13**). This showed that the chance of finding long TEs (i.e., >1,000 bp) of the same family in two randomly drawn genomic regions is low—for instance, within 1,000 randomly distributed region pairs of 100 kb in chromosomes 1 and 7—we expect to find on average less than 0.2 TEs of the same family (**Supplementary Figure S13a)**. Conversely, we find insertions of specific TE-families (**Figure 2**) longer than 1,000 bp in all of the breakpoints on chromosomes 1 and 7 in NEAC, strengthening the association between TEs and these inversion breakpoints. When considering short family-specific TEs (i.e. below 1,000 bp), we did not observe any deviation from the average TE count in random regions (**Supplementary Figure S13**).

### Accumulation of hAT-transposons in chromosome 1 and 7 breakpoints: Evidence for tandem MITE insertions

The breakpoints of chromosomes 1 and 7 in NEAC harbored insertions of the same two particular families initially annotated as DNA *hAT* transposons. This is a common TE-superfamily characterized by the presence of TIRs flanking an open reading frame (ORF) that encodes a transposase in their autonomous form (Atkinson, 2015). These two breakpoint families were termed gadMor-*hAT*-1784 and gadMor-*hAT*-8043 (**Figure 2-3a**). The gadMor-*hAT*-8043 insertions had a mean length of 1,973 bp (5 TEs; *SD*: 454 bp) and 5,838 bp (2 TEs) in chromosomes 1 and 7, respectively, whereas gadMor-*hAT*-1784 had a mean length of 1,334 bp (3 TEs; *SD:* 84 bp) in chromosome 1, and two insertions in the chromosome 7 breakpoint A (mean length 1,404 bp), overlapping an insertion of the gadMor-*hAT*-8043 family (**Figure 3a**). In the syntenic breakpoint regions in NCC, gadMor-*hAT*-1784 and gadMor-*hAT*-8043 were present only in the syntenic breakpoint A in chromosome 1 (mean lengths: 1,385 bp and 890 bp for gadMor-*hAT*- 1784 and gadMor-*hAT*-8043, respectively).

**Figure 3:**
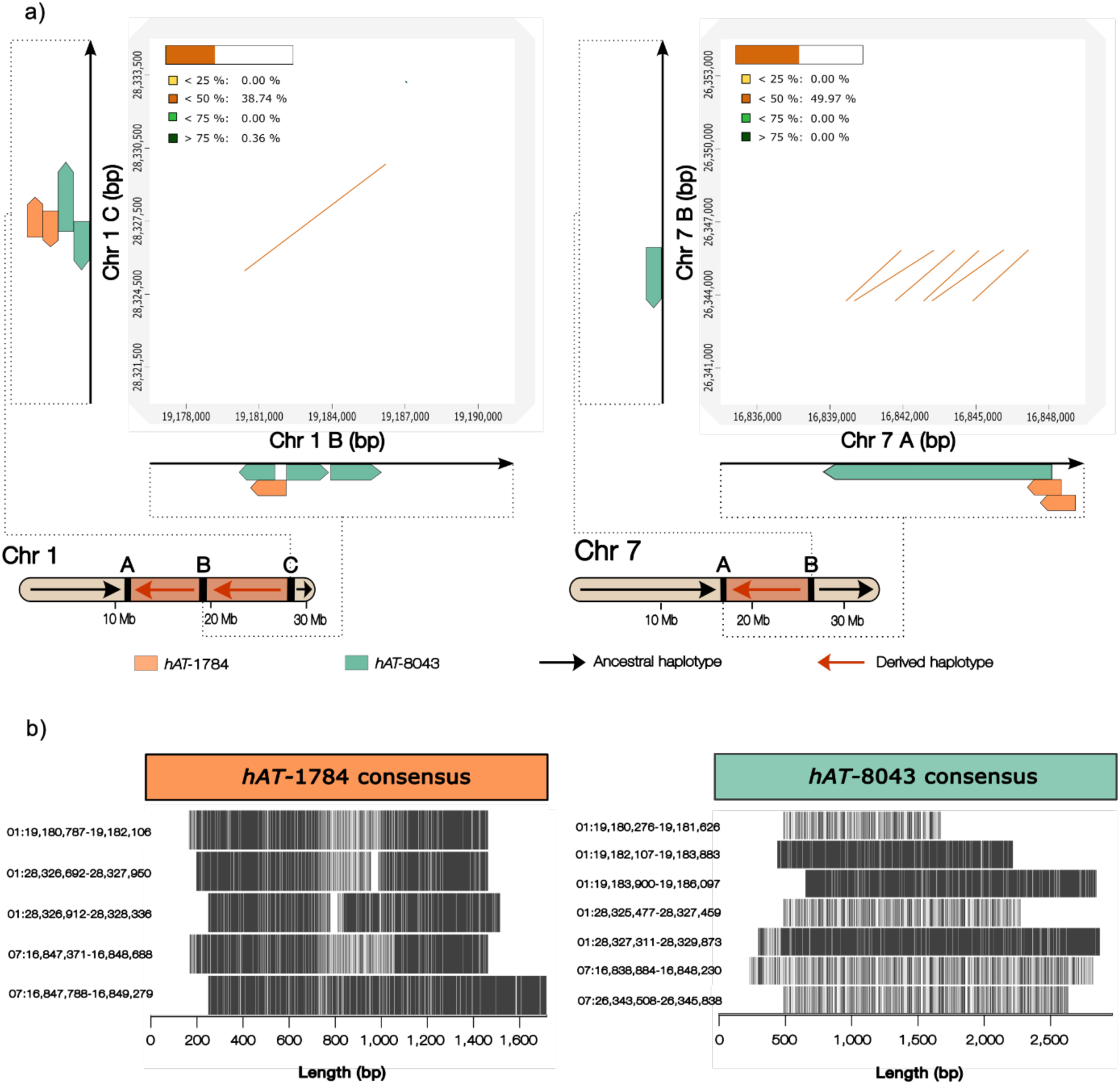
Accumulation of *hAT* TEs in the inversion breakpoints on chromosomes 1 and 7 in NEAC. **a,** Dot-plot alignments of the breakpoint pairs in the derived (inverted) haplotypes on chromosome 1 (breakpoint B vs. C) and chromosome 7 (breakpoint A vs. B), highlighting sequence similarity between breakpoint regions (full alignments are shown in **Supplementary Figure S11**). Percentage identity of alignments are colored according to the inset and correspond to regions with insertions from two *hAT* families, gadMor-*hAT*-1784 (orange) and gadMor-*hAT*-8043 (green). Insertions are drawn to scale, with strand orientations indicated on both axes. Red arrows denote the derived (inverted) haplotype. Chromosomes are drawn to scale, with sizes indicated in Mb. **b**, Mapping of the breakpoint insertions of gadMor-*hAT-*1784 and gadMor-*hAT-*8043 onto their respective consensus sequences, illustrating the portion of each consensus represented by the different copies. Sequence identity with the consensus is shown by black lines.

The insertions of gadMor-*hAT*-1784 elements were either overlapping the center of the breakpoints (e.g., breakpoint B and C in chromosome 1) or located in the immediate vicinity of the breakpoints (chromosome 7). Moreover, the breakpoint regions containing these *hAT* insertions exhibited a tandem repetitive pattern when aligned (**Figure 3a**), and the inserted *hAT* copies represented the same part of the gadMor-*hAT*-1784 consensus sequence (**Figure 3a-b**; **Supplementary Table S6**). We observed the same patterns for the gadMor-*hAT*-8043 breakpoint insertions, which had overlapping insertions with several of the gadMor-*hAT*-1784 insertions (**Figure 3b**).

To investigate the structural characteristics of gadMor-*hAT-*1784 on chromosomes 1 and 7, we utilized TE-Aid (Goubert et al., 2022). Self-alignment dot-plots of the gadMor-*hAT-*1784 consensus sequence revealed a complex repeated sequence motif consisting of two large TIRs of ∼600 bp, which were made of multiple tandem repeats with repeating units of 167-169 bp (**Figure 4a-b**). We did not identify any ORFs in the consensus sequence of gadMor-*hAT*-1784, and mapping of the consensus to the genome showed high genomic coverage for the long 600 bp TIRs, but not for the short internal sequence of the consensus (**Figure 4c**). This indicates a near-complete loss of the internal transposase-encoding sequence of the gadMor-*hAT*-1784 TE. The presence of inverted repeats and internal deletions suggests that gadMor-*hAT-*1784 is a *hAT*-derived non-autonomous MITE with multiple internally inverted repeats in tandem (**Figure 4a-b**).

**Figure 4:**
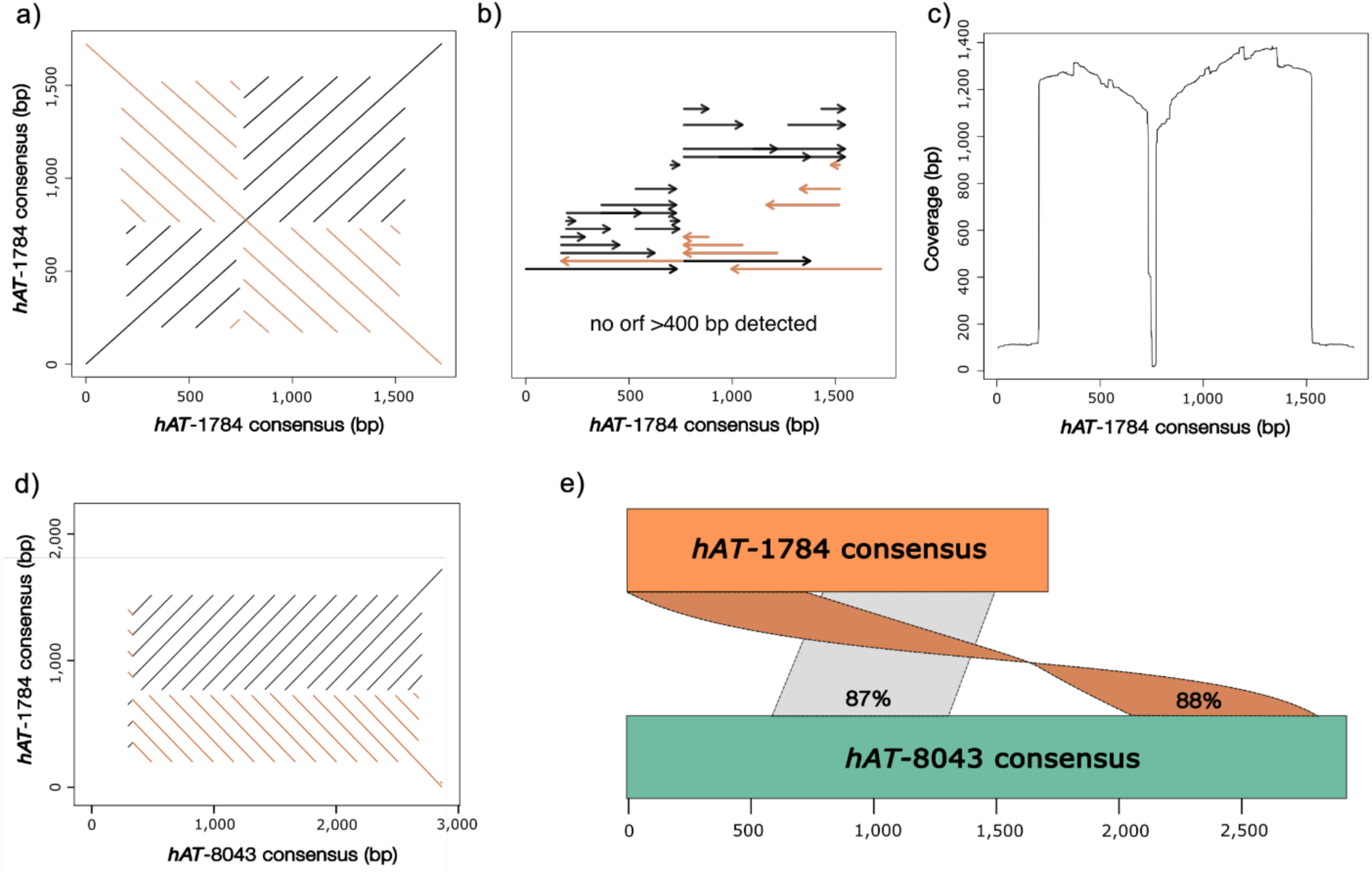
Structural characteristics of the *hAT*-derived MITEs in Atlantic cod. **a,** Self-alignment dot-plot of the gadMor-*hAT*-1784 consensus sequence, showing the inverted repeat structure. **b,** Internal repeats of the gadMor-*hAT-*1784 consensus, with black arrows indicating forward repeats and orange arrows indicating inverted repeats. **c,** Genomic coverage of the gadMor-*hAT*-1784 consensus, showing the pileup of genomic hits across the consensus sequence. **d,** Dot-plot alignment between the gadMor-*hAT*-1784 and gadMor-*hAT*-8043 consensus sequences, showing sequence similarity between the two TE families. **e,** Mapping of the gadMor-*hAT*-1784 (orange) onto the gadMor-*hAT*-8043 (green) consensus, with sequence identity for mapped regions indicated in percentages. Alignments in the forward direction are shown in gray, while reverse orientations are shown in orange.

We performed the same TE structural analysis on the gadMor-*hAT-*8043 consensus, which yielded a repetitive pattern resembling a simple sequence repeat with similar repeating units of ∼167 bp (**Supplementary Figure S14**). Since the gadMor-*hAT*-1784 and gadMor-*hAT*-8043 families shared a tendency to overlap the breakpoints in chromosome 1 and 7, and were often annotated in the vicinity of each other in a tandem arrangement of similar sized repeating units, we tested whether they share a common origin. We therefore compared the consensus sequences and structural characteristics of the two families following the 80-80-80 rule (Wicker et al., 2007) for families and 80-95-98 rule for subfamilies. This revealed that gadMor-*hAT*-1784 and gadMor- *hAT*-8043 are two different non-autonomous subfamilies of the same ancestral *hAT* family (**Figure 4d-e**).

We validated the repetitive structure of both *hAT* subfamilies by performing self-alignment dot-plots of the breakpoint regions spanning the insertions of the gadMor-*hAT*-1784-derived MITE and gadMor-*hAT*-8043-derived simple sequence repeat on chromosomes 1 and 7, confirming the presence of long tandemly inserted MITE and simple sequence repeats within the breakpoints, which resemble the same parts of the gadMor-*hAT*-1784 and gadMor-*hAT*-8043 consensus sequences (**Figure 5**).

**Figure 5:**
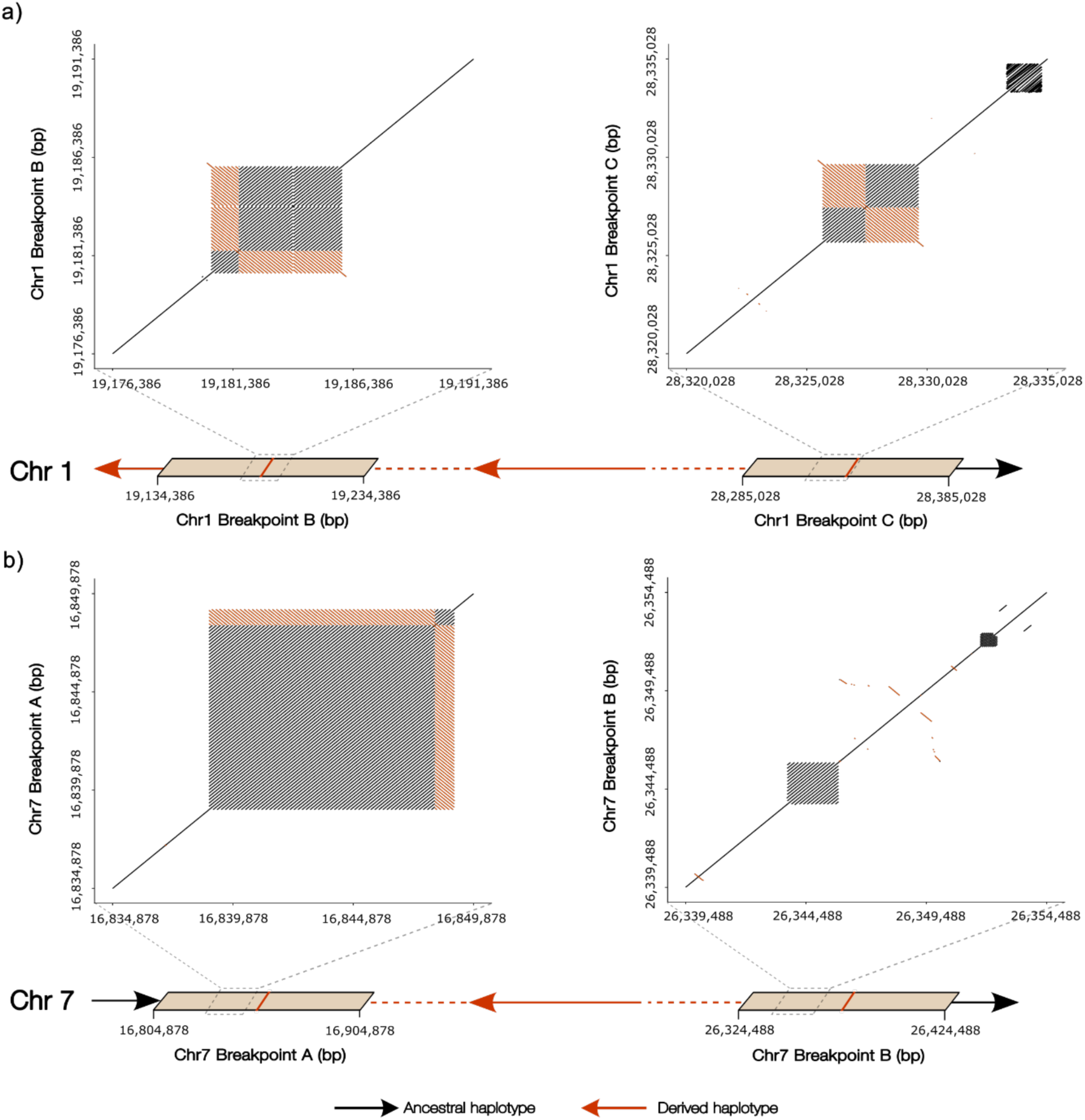
Self-alignment dot-plots of breakpoint regions on chromosomes 1 and 7 containing gadMor-*hAT*-1784 and gadMor-*hAT*-8043 insertions. **a,** Chromosome 1 breakpoint self-alignment. **b,** Chromosome 7 breakpoint self-alignment. Red arrows indicate an inversion, and black arrows indicate regions outside the inversion. The red lines represent the midpoint coordinates of the breakpoint estimates from **Table 1**. Alignments in the forward direction are shown in gray, while reverse orientations are shown in orange.

### Inversion breakpoints on chromosome 2 contain heterogenous TE-family landscapes

We identified insertions from several TE families within the breakpoints of the chromosome 2 inversion (**Figure 1**). These families represented a diverse landscape of TE families, often with multiple insertions (2-5 per family) of varying length (131-1,757 bp), primarily belonging to the retroelements. Most of these TE insertions were shared between the cod ecotypes (NEAC and NCC). However, two LTR families—gadMor-Ty3/Metaviridae-342 and gadMor- Ty3/Metaviridae-93—were specific to the derived breakpoint regions in NCC, and absent from the ancestral breakpoint regions in NEAC.

To investigate the breakpoints in more detail and evaluate the TE insertion patterns across individuals (and not only in the reference genomes), we used long range PCR and long read sequencing (PacBio HiFi; see **Materials and Methods**) to amplify the chromosome 2 breakpoint regions. We successfully sequenced 39 individuals of NEAC and NCC from different locations (**Supplementary Table S7**), including homozygous and heterozygous individuals for the two inversion haplotypes. By mapping reads from heterozygous individuals to the NEAC and NCC reference genomes (using coordinates from **Table 1**), we were able to validate our bioinformatics-led estimations of the breakpoint coordinates, and further refine the breakpoint intervals on chromosome 2 to ranges of 111-266 bp (**Supplementary Table S8** and **Supplementary Figure S15**).

One notable feature of these breakpoint sequences was their low sequence variation across the sequenced individuals (**Supplementary Figure S15**). This pattern extended to the level of TE insertions, where one of the two LTR families—gadMor-Ty3/Metaviridae-342—was consistently found in the breakpoint regions of all individuals carrying the inverted haplotype. These insertions represented internal, protein-coding regions of the TE family (**Supplementary Table S6**), and the insertions in breakpoint A were oriented in the reverse direction of the breakpoint B insertions. The regions harboring the remaining TE breakpoint families that were shared across NEAC and NCC were not successfully amplified, as they were located outside the range of the sequenced regions.

## Discussion

We have identified and characterized the TE content in the breakpoint regions for the four chromosomal inversions previously identified in Atlantic cod (**Table 1**, **Figure 2**) (Berg et al., 2016, 2017; Matschiner et al., 2022). This involved obtaining a curated lineage-specific TE library **(Supplementary Table S2-S3** and **Supplementary Figure S5-S6)**, as suggested by best practices for proper characterization and annotation of repetitive DNA content (Platt et al., 2016; Peona et al., 2021; Goubert et al., 2022; Storer et al., 2022). Our most notable finding is that certain TE orders (e.g., MITEs and Mavericks) and superfamilies (*hAT*-elements) display elevated densities (i.e., the proportion of TE insertions per sliding window) within breakpoints relative to other areas of the genome (**Figure 1** and **Supplementary Figure S10**). This association between TEs and breakpoints is largely explained by the presence of the same TE families in breakpoint pairs, illustrated by the accumulations of the two *hAT*-derived MITE subfamilies in the breakpoints of the derived variants (i.e. the inverted state) in chromosomes 1 and 7 in NEAC (see **Figures 3 and 5**).

We found that the breakpoint pairs of all four inversions contained family-specific TE accumulations (**Figure 1**). Specifically, we found that the breakpoints of the derived haplotypes for chromosomes 1 and 7 harbored long, related TEs (>1,000 bp), which we demonstrate is an unlikely association to observe by chance across the genome (**Supplementary Figure S13**). When aligning the two breakpoint regions associated with each inversion, we found a higher sequence identity in regions that contained related TEs compared to adjacent regions (**Supplementary Figures S11**). Furthermore, our amplification of the breakpoint regions on chromosome 2 in multiple cod individuals showed that the breakpoint insertions of TEs were conserved across the sequenced individuals, which can be extrapolated to the remaining three inversions in chromosomes 1, 7 and 12. Together, the observation of sequence similarity and preserved insertions of specific TE families seems to represent a general feature of the four major inversions in the Atlantic cod.

We observed tandemly repeated *hAT-*insertions within the inversion breakpoints of chromosomes 1 and 7 (see **Figures 3 and 5**). These tandem insertions were identified using a combination of automated approaches (Flynn et al., 2020; Orozco-Arias et al., 2024) and manual curation (Goubert et al., 2022) (see **Supplementary Figure S5**), including the validation of the repeated structure of the breakpoint insertions with self-alignment dot-plots (**Figures 4-5**). *hAT*- elements are known to be abundant in teleost fish (Gao et al., 2016), and previous studies have suggested *hAT*s as drivers for creating inversions in fruit fly (*Drosophila melanogaster*) (Lyttle & Haymer, 1992), maize (*Zea mays*) (Zhang & Peterson, 2004; Sharma & Peterson, 2023), yeast (*Saccharomyces cerevisiae*) (Sarilar et al., 2015), and structural variants observed in aphids (Mathers et al., 2021). We classified the breakpoint *hAT* elements in the Atlantic cod as MITEs because they lacked the transposase but still preserved the inverted repeat regions. This is particularly evident from the gadMor-*hAT*-1784 consensus sequence, which lacks a transposase **(Figure 4)**, and is further exaggerated in the subfamily gadMor-*hAT*-8043, which has only preserved one of the TIRs in a tandem repeat arrangement. The complex, nested pattern of *hAT*- derived MITEs (see **Figures 3 and 5**) is unique to the breakpoint pairs of chromosomes 1 and 7, suggesting a putative causal link between these elements and the origin of the two inversions.

The presence of long inverted repeats likely facilitated the origin of inversions through ectopic recombination. Inversions that occur by non-homologous end joining (NHEJ) leave distinct genomic signatures that are independent of TE accumulations (e.g., inverted duplications of genes and non-repetitive DNA) (Ranz et al., 2007; Guillén & Ruiz, 2012), whereas inversions formed by ectopic recombination are typically characterized by accumulations of related TEs and inverted repeats (Delprat et al., 2009; Guillén & Ruiz, 2012). Therefore, one way to distinguish these mechanisms is by investigating the TE sequence similarity and elucidating the shared structural characteristics (e.g., TIRs and LTRs) of their insertions – as done here. In line with an ectopic recombination scenario, the characterization of the breakpoint sequences revealed long stretches of inverted repeats (**Figure 5**), in congruence with long MITE insertions. Indeed, MITEs are associated with inversions in various study systems, such as *Drosophila repleta* (Rius et al., 2013) and *Brassica napus* (Wan et al., 2017), and recombination between short inverted sequences of TEs have been shown to generate inversions in bacterial chromosomes (Ling & Cordaux, 2010). Our results demonstrated accumulations of *hAT*-derived MITEs spanning several thousands of base pairs in the breakpoints of chromosomes 1 and 7 (mean lengths spanning 1,334-5,838 bp, **Figure 3**) with long repeating units (167-169 bp). We propose that the presence of these long inverted tandem repeats facilitated the inversions by ectopic recombination. This scenario is further supported by our manual inspection of the two *hAT*-consensus sequences (gadMor-*hAT*- 1784 and gadMor-*hAT*-8043) and their breakpoint insertions, which confirmed that their insertions consistently represented the same or overlapping portions of their respective consensus sequences (**Supplementary Table S6**).

Our results for chromosomes 1, 7 and 12 revealed a distinct pattern where certain related TEs only associated with the breakpoints of the derived haplotypes, while being absent in the corresponding homologous regions in the ancestral arrangement (**Figure 2**). The TEs found on these three chromosomes belonged primarily to the DNA class II elements containing TIRs. Upon activation of a transposase (either autonomously or non-autonomously), these TEs may undergo expansions that leave both their insertion sites and site of origin susceptible to structural alterations (Sharma & Peterson, 2023). Thus, the presence of multiple DNA elements in the derived breakpoints of chromosomes 1, 7 and 12, corroborates with previous studies on the role of class II TE activity in driving structural variants, including inversions in *Zea mays* (Zhang & Peterson, 2004; Sharma et al., 2021; Sharma & Peterson, 2023) and *Drosophila melanogaster* (Lyttle & Haymer, 1992).

In contrast, our analysis of the chromosome 2 breakpoints revealed a predominance of retroelement insertions, many of which were present in both the ancestral and derived breakpoints (**Figure 2**). Chromosomal rearrangements in certain avian genomes (Boman et al., 2019) and yeast (*Saccharomyces cerevisiae*) (Chan & Kolodner, 2011) show a positive correlation with enrichments of retroelements, particularly the LTRs. Unlike DNA elements, retroelements transpose via reverse transcription, which leaves behind copies at their original genomic locations. Thus, by comparing the ancestral and derived states of the breakpoint regions, we can identify signatures left by different TE classes that might offer insights into their potential involvement in the origins of the inversions.

However, in the case of the HiFi-sequenced breakpoints on chromosome 2, we only sequenced regions harboring one of the LTR families that were associated with the derived and not the ancestral breakpoint regions (i.e., gadMor-Ty3/Metaviridae-342). Common for these breakpoint insertions was their location at the outer edges of the two breakpoint regions, with the different breakpoints harboring insertions in the opposite orientation of one another. Each breakpoint insertion represented internal parts (ORFs) of their consensus sequences. Furthermore, the insertions showed substantial divergence from their consensus sequences, with a mean divergence of 31.1% for the gadMor-Ty3/Metaviridae-342 insertions (**Supplementary Table S6**). The high divergence suggests that the breakpoint insertions represent old, non-autonomous TE copies that may have been preserved at these sites since before the origin of the inversion, which is estimated at 0.88 Mya (Matschiner et al., 2022).

The identification of TE-derived tandem repeats within the various inversion breakpoints adds to recent discoveries of the impact of tandem-repeated TEs in shaping genomes. This is exemplified in the 9 kb tandemly inserted TE influencing melanism in the peppered moth (Hof et al., 2016). Inserted TEs can elongate by unequal crossover and replication slippage, or they can be inserted in tandem configurations (McGurk & Barbash, 2018). Indeed, DNA class II transposons, such as the ones found here in the inversion breakpoints, commonly perform double insertions in a head-to-tail orientation (McGurk & Barbash, 2018), making them susceptible for ectopic recombination and subsequent structural alterations, such as inversions. Given the extensive TE diversity and particular enrichment of class II TEs in teleost fish (Gao et al., 2016; Reinar et al., 2023), along with the prevalence of inversions (Natri et al., 2019; Matschiner et al., 2022; Andersson et al., 2023; MacGuigan et al., 2023) and other structural variants documented in cod and other teleosts, e.g., loss of MHC II in cod (Star et al., 2011; Malmstrøm et al., 2016), loss of MHC II in pipefishes (Haase et al., 2013; Small et al., 2016; Roth et al., 2020), loss of MHC II in anglerfish species (Dubin et al., 2019; Swann et al., 2020), hemoglobin MN cluster duplications in nine-spined stickleback and codfishes (Baalsrud et al., 2017; Varadharajan et al., 2019), and copy number variations of antifreeze glycoprotein-encoding genes in codfishes and notothenioids (Baalsrud et al., 2018), it becomes essential to thoroughly characterize and quantify TEs as potential evolutionary drivers of structural variants in general. Only then will we be able to obtain a better understanding of the drivers of the origin of structural variants. Our study highlights the propensity of certain TE structural features, such as TIRs, to promote inversions when present in tandem arrangements. In contrast to inversions originating from NHEJ, all of the large inversions on chromosome 1, 2, 7 and 12 in Atlantic cod likely originated through ectopic recombination involving TEs, both DNA elements and retroelements. Furthermore, the conserved breakpoint positions and breakpoint-specific TE insertions across individuals support separate ancient events for each inversion predating the western and eastern Atlantic cod population split in line with the phylogenetic dating analyses of Matschiner et al., (2022).

## Materials and Methods

### Data

We used three high-quality reference assemblies for this work: the Northeast Arctic Cod (NEAC) assembly (NCBI accession ID: GCF_902167405.1; contig N50 = 1,015 kb, scaffold N50 = 29 Mb, and BUSCO score = 96.8 %), the Norwegian Coastal Cod (NCC) genome assembly (NCBI accession ID: GCA_964260575; contig N50 = 271 kb, scaffold N50 = 28 Mb and BUSCO score = 90.5 %), which are both based on PacBio long read sequencing and chromosome-resolved with Hi-C linked reads; and the Oxford Nanopore-based Celtic cod assembly (NCBI accession ID: GCA_010882105.1; contig N50 = 10,560 kb, scaffold N50 = 27 Mb, and BUSCO score = 94.1 %). Detailed description on sequencing and assembly construction of the three assemblies can be found in Jentoft et al. (2024), Hoff et al. (2024), and Kirubakaran et al. (2020), respectively.

### Estimating the inversion breakpoint coordinates

To determine the location of the breakpoint regions for the inversions located on chromosomes 1, 2, 7 and 12, we aligned the reference genome for NCC to the reference genome of NEAC using SyRi (Goel et al., 2019). This allowed the identification of syntenic regions and structural variants between the two reference genomes, including inversions. As a complementary analysis, we dot-plotted chromosomes 1, 2, 7 and 12 from NEAC against their corresponding homologous chromosomes in NCC to visually inspect the presence of inversions with D-GENIES v.1.4 (Cabanettes & Klopp, 2018). The breakpoint regions were not identified as exact locations on either genome, but rather as approximate regions in which the breakpoints were located (**Supplementary Table S1a; Supplementary Figures S1-S2**), surrounded by clear homologous regions between the two genomes, one of which was in opposite direction to the other. We cross-referenced the breakpoint coordinates with the inversion configurations and breakpoints present in the Celtic cod assembly, which is assumed to have inversion haplotypes that are similarly oriented as the NCC (Kirubakaran et al., 2020). Our synteny-based identification of breakpoints indicated that the chromosome 7 and 12 inversions might be double inversions, the former with overlapping breakpoints. Since we do not have population data to support this (nor do any other previous contributions on the cod inversions clearly suggest this), we are treating them as single inversions (breakpoint locations highlighted in **Supplementary Figure S2**).

To obtain more precise estimations of the breakpoint regions, we mapped raw reads from the NEAC and NCC assemblies back onto both of the assemblies using minimap v.2.26 (Li, 2018). When aligning reads back to their original assembly (e.g., NEAC reads to the NEAC reference genome), reads that spanned an inversion breakpoint mapped continuously, without interruption. In contrast, when reads were aligned to the alternative assembly (e.g., NEAC reads to the NCC reference genome), those spanning the inversion breakpoints would only partially map to one boundary, with the remaining portion aligning in reverse orientation to the opposite boundary. We evaluated read coverage around putative breakpoint regions using pyGenomeTracks (Lopez-Delisle et al., 2021) (**Supplementary Figure S3**), where we assumed that there would be a lower coverage around breakpoint regions from cross-mapped reads (e.g., from NCC onto NEAC) than from mapping reads from the same assembly (e.g., NEAC onto NEAC). Finally, we visually inspected hard-clipped reads (i.e. reads where a portion has not aligned well to the reference genome well and is thus excluded from the alignment) as labeled by minimap 2.26, using GenomeRibbon (Nattestad et al., 2021). The hard-clipped reads that mapped to two locations in opposite directions (i.e., mapped to two breakpoints) marked the boundaries of the inversion breakpoint intervals (**Supplementary Figure S4**). To infer syntenic breakpoint regions in the ancestral (non-inverted) haplotypes, we used the estimated breakpoint intervals and adjacent regions (midpoint of breakpoint interval ± 50,000 bp) and mapped these regions onto the assembly carrying the ancestral haplotype (for each inversion), using minimap v.2.26. The outer boundaries of the mapped regions defined the edges of the syntenic breakpoint regions (**Supplementary Table S1b**).

### HiFi-sequencing and population-level variant calling of chromosome 2 breakpoints

We obtained 58 tissue samples from two wild populations of Atlantic cod, Lofoten (N=30) and Averøya (N=28). Additionally, we obtained three individuals from the Atlantic cod breeding program for aquaculture at Nofima, Tromsø, for establishing and optimizing the PCR protocol for sequencing the inverted and non-inverted breakpoint region in chromosome 2. DNA was extracted from 0.025g tissue samples using the DNeasy Blood and tissue mini kit by QIAGEN. All samples were extracted according to the manufacturer’s instructions in “DNeasy® Blood & Tissue Handbook, July 2020”. The concentration and purity of the DNA was estimated using NanoDrop (Thermo Fisher Scientific) and a Qubit fluorometer (Invitrogen, Thermo Fisher Scientific).

Library preparations for PacBio HiFi sequencing were done by the Norwegian sequencing center (NSC) on ∼25% of 8M SMRT cells using Sequel II Binding kit 2.0 and Sequencing chemistry v2.0. The sequencing data was demultiplexed with the Demultiplexing pipeline on SMRT Link v10.2.0.1333434, and circular consensus sequencing (CCS) reads were then generated To analyze breakpoint variation in chromosome 2 we mapped sequenced against NEAC and NCC reference genomes using minimap2 (Li, 2018). We were able to map a total of 158 CCS sequences, and we used SAMtools (Danecek et al., 2021) to examine the mapped output bam files, and BCFtools mpileup to call variants (Danecek et al., 2021) for NEAC and NCC. The resulting VCF files were used for haplotype phasing and tagging with WhatsHap v1.4 (Martin et al., 2016). These haplotypes were then assembled using Flye v2.9 (Kolmogorov et al., 2020). We then created *de novo* haplotype assemblies that we aligned together in a multiple sequence alignment using Geneious Prime 2021.2 (https://www.geneious.com) (**Supplementary Figure S15**).

To determine the breakpoint regions on chromosome 2 and their sequence variation, we compared the breakpoint sequences of each individual to bioinformatically determined breakpoint regions on the NEAC and NCC genome assemblies (**Table 1**). We focused on heterozygous individuals and compared the derived (inverted) breakpoint A to the syntenic breakpoint A region of the ancestral (non-inverted) haplotype (**Supplementary Figure S15**), and performed the comparison for the breakpoint B haplotypes (**Supplementary Figure S15**).

### Repeat annotation and classification

To capture species-specific TE composition in Atlantic cod, we started by producing a *de novo* repeat library with RepeatModeler2 (Flynn et al., 2020). We ran RepeatModeler v2.0.1 on the NEAC assembly and included the built-in LTR structural discovery pipeline (Ellinghaus et al., 2008; Ou et al., 2018) to increase LTR detection. We then used MCHelper (Orozco-Arias et al., 2024) for automatic curation of our TE library, which involves identifying and removing false positives (i.e., multi-copy genes, short tandem repeats, etc.), redundancy reduction, TE consensus elongation, structural checking using TE-Aid and TE classification as suggested by best practices (see Goubert et al., 2022). MCHelper was run with the fully automated setting, with simple-sequence repeat detection set to 90% and consensus Blast, Extract and Extension (BEE) cycle set to 8 rounds in MCHelper. We then focused on the non-redundant, unclassified TEs for manual processing and curating TEs. An overview of the pipeline we constructed for manual curation of TEs can be found in **Supplementary Figure S5**. In short, the pipeline consists of four steps: (1) TE classification of the non-redundant library using RepeatClassifier v.2.0.4 (only the Dfam 3.8 curated database) and PASTEC (Hoede et al., 2014); (2) generation of a priority list for manual curation by masking the NEAC assembly with the unclassified, non-redundant TE library using RepeatMasker v.4.1.5 (Smit et al., 2013). Priority was based on insertions in the vicinity of inversion breakpoints and general abundance in the genome; (3) Manual curation of prioritized TEs using the output from PASTEC (structural and homology-based evidence), RepeatClassifier (built-in homology-based tool in RepeatModeler2), MCHelper, and manually inspecting the output from TE-Aid and multiple sequence alignments of TE consensus aligned to the assembly. Finally, we performed (4) automatic curation of the remaining uncurated library using RMBlast v.2.13.0 and the curated TEs as a database. This outputted a final curated, classified TE library that was used to mask the NEAC, NCC and Celtic cod assemblies using RepeatMasker v.4.1.5 with slow search for high sensitivity searches (see library comparisons in **Supplementary Figure S16**). To evaluate the quality of the final TE library, we compared the length and contiguity of the TE annotations of the raw vs. the final curated library using the NTE50 and LTE50 metrics, as suggested by Jamilloux et al. (2016). NTE50 represents the number of largest TE copies required to annotate 50% of repetitive DNA, while LTE50 refers to the length threshold at which 50% of the repetitive DNA is annotated by copies longer than this length. Furthermore, the final TE library contained an overall longer and fewer consensus sequences (**Supplementary Figure S16**), indicating less false positives, lower consensus redundancy, and more complete consensus sequences.

To determine the population-level frequencies of TEs in breakpoints, we used the HiFi-reads of the inversion in chromosome 2 and examined whether these reads overlapped with positions of annotated TEs using a genome browser (IGV v.2.12.3). We also used the Atlantic cod species-specific TE-library to mask individual reads to confirm each observed breakpoint TE.

### Aligning inversion breakpoints to investigate the presence of related TEs

To determine the repeat landscape of the different breakpoints and better capture the TE content in the regions flanking a breakpoint in a broader genomic context, we extended each breakpoint interval (of the derived inversion haplotypes) by taking the interval midpoints and expanding the regions by 50 kb in both directions. This resulted in equally sized breakpoint regions of 100 kb.

We evaluated the sequence similarity between breakpoint pairs of the four inversions by dot-plotting the breakpoint region pairs associated with each inversion using D-GENIES v.1.4 (Cabanettes & Klopp, 2018). If TEs are associated with an inversion – in concordance with a scenario of ectopic recombination; we expect the breakpoints to align due to similarity of repeats. We also expect to find related TE copies within the aligned regions. To ensure we limited our investigation to putative homologous sequences, only alignments with sequence identity exceeding 25% were retained and screened for related TEs. TE fragments that were less than 100 bp in length, and thus considered less likely to facilitate ectopic recombination, were excluded, and the remaining TEs were grouped by family. The TE families that were present in breakpoint pairs were manually investigated by curation of the consensus with structural and homology-based evidence from PASTEC (Hoede et al., 2014), MCHelper (Orozco-Arias et al., 2024), and manual searches in the Dfam (Storer et al., 2021) and Censor (Kohany et al., 2006) databases, and by evaluating the insertions of copies to evaluate the number and length of the insertions, mean divergence from the respective consensus sequences, and whether the insertions overlapped the same parts of the consensus.

### Structural validation of the gadMor-hAT-1784-derived MITE

We used TE-aid to validate structural features of MITEs associated with gadMor-*hAT-* 1784 that appeared in multiple inversions (Goubert et al., 2022). Given the presence of numerous internally inverted repeats within the consensus sequence of gadMor-*hAT-*1784, we extracted the breakpoint regions from chromosomes 1 and 7 containing gadMor-*hAT-*1784 insertions, and used self dot-plots of each breakpoint to visualize palindromic insertions. To obtain this dot-plot we used BLASTn v.2.6.0+ (Altschul et al., 1990; Camacho et al., 2009) to compare the breakpoint insertions to the palindromic gadMor-*hAT-*1784 consensus.

We aligned the gadMor-*hAT*-1784 consensus sequence with the overlapping gadMor-Unc- 8043 consensus with MAFFT v.7.520 (L-INS-i) (Katoh & Standley, 2013)to investigate sequence similarity. To further evaluate if the two consensus sequences were part of the same family and/or subfamilies, we used CD-HIT v.4.8.1 (Fu et al., 2012) and applied the 80-80-80 rule for family identification (i.e., family members are at least 80 nucleotides in length, and share 80% of residues over at least 80% of their length), and the more stringent 80-95-98 rule for subfamily identification.

This showed that gadMor-Unc-8043 and gadMor-*hAT*-1784 are two subfamilies of the same family, and the two consensus sequences were grouped together as the same *hAT* family (gadMor- *hAT*-1784) for further analyses.

Additionally, to confirm the tandem-repeated MITE-structure inherent in the gadMor-*hAT-* 1784 insertions, we employed ULTRA (Olson & Wheeler, 2018) to identify and annotate tandem repeats in the NEAC assembly. ULTRA v.0.99.17 was run with a maximal detectable repeat period of 1,000 bp, and we checked whether the annotated tandem repeats overlapped with the gadMor-*hAT-*1784 and gadMor-*hAT*-8043 insertions in the chromosome 1 and 7 inversions in NEAC using a genome browser (IGV v.2.12.3).

### Estimating putative centromeres

To obtain statistical inferences of the density of TEs in breakpoint regions (see section *Statistical assessment of TE features within breakpoints* below), we needed to account for centromeric regions, as these regions are comprised of high density repeats, potentially biasing our inferences. We predicted putative centromeric regions *in silico* using three independent approaches. First, we plotted the density of interspersed repeats (from our TE library) and tandem repeats (as annotated by the TRF module included in RepeatMasker) on chromosomes 1, 2, 7 and 12, using non-overlapping sliding windows of 10 kb. Second, we downloaded the putative 158 bp centromeric repeat sequence that is suggested to represent the centromeric repeat in Celtic cod (Kirubakaran et al., 2020), and mapped the sequence onto chromosomes 1, 2, 7 and 12 in the NEAC and NCC assemblies using BLAST v.2.14.1 (Altschul et al., 1990; Camacho et al., 2009). We compared the density peaks of interspersed and tandem repeats with the locations of the Celtic cod centromere sequence, and used the corresponding density peaks for further processing. Third, we used CentroMiner v.1.2.1 (Lin et al., 2023) to predict centromere candidates on each of the four chromosomes, and compared the output to the general repeat density peaks and the Celtic cod centromere sequence. Finally, we extracted the regions containing candidate centromeric regions and self-aligned these regions to identify potential repetitive motifs (e.g., simple repeats and palindromes), characteristic of centromeres. This holistic approach provided a final set of coordinates for the putative centromeres in these four chromosomes, which were excluded in further statistical analyses.

### Statistical assessment of TE features within breakpoints

We conducted two distinct statistical analyses on all inversion breakpoints to determine TE density and the probability of finding similar TE copies in breakpoint pairs. First, we plotted the mean TE density across chromosomes 1, 2, 7 and 12 in NEAC and NCC using non-overlapping sliding windows of 25 kb, 50 kb, and 100 kb. We present the results using the 50 kb windows, since it was coherent with the size of the flanking sides of the breakpoint regions. To test whether the TE density in the breakpoints was significantly different from the chromosomal means, we did a Student’s t-test for each chromosome, excluding centromere regions as described above. We repeated this analysis at the level of TE orders and superfamilies. We used a t-test for comparison because this test compares means and not median values, which would introduce bias in the results due to the density estimates being skewed towards zero in many windows, especially at the lower taxonomic levels of TEs. Additionally, we checked if the t-tests were biased due to the priority of TEs in the curation process (**Supplementary Figure S17**), by applying the same test to the raw, uncurated TE library. This showed a higher detection of MITEs after curation, but our results were unaffected at the level of superfamilies (e.g., the high levels of *hAT*-elements in chromosome 7) (**Supplementary Figure S17**).

Second, we assessed the probability of encountering copies of the same TE family within both breakpoints of an inversion. For chromosomes where we found inversions (1, 2, 7 and 12), we generated 1,000 randomly selected region pairs (mock breakpoints) along the chromosome, with each region being 10 kb in size. We then calculated the average count of related TEs (from the same family) in the mock breakpoints, using different minimum TE insertion lengths: 0 bp, 100 bp, 500 bp, 1,000 bp, and 2,000 bp. This allowed us to estimate the probability of encountering longer TE insertions from the same family in randomly selected region pairs. We repeated this analysis for increasing region sizes, ranging from 10 to 100 kb in 10 kb increments. The average counts of related TEs in the mock breakpoints were then compared to the observed counts of related TEs in the real inversion breakpoints.

## Acknowledgements

This work was funded by the Research Council of Norway through the following projects: **‘Nansen Legacy’** (RCN no. 276730), **‘EBP-Nor’** (RCN no. 326819), **‘Comparacod’** (RCN no. 222378) and the University of Oslo. We thank Ave Tooming-Klunderud, Norwegian Sequencing Centre (NSC), University of Oslo for assistance with PacBio sequencing of Atlantic cod individuals. We also thank Dr. Johann Confais, Unité de Recherche Génomique Info (URGI) for assistance with TE classification, Prof. Dr. Alexander Suh, University of Bonn for input on analyses, and in particular Dr. Simon Orozco-Arias, Centre for Research in Agricultural Genomics (CRAG) for help and discussion of TE curation. All sequencing library creation and high throughput sequencing was carried out at the NSC, University of Oslo, Norway. All computational work was done on the Saga Supercomputing Cluster (Norwegian metacenter for High Performance Computing, NOTUR, and the University of Oslo) operated by the Research Computing Services group at USIT, the University of Oslo IT-department (http://www.hpc.uio.no/).

## Data availability

Data is available in the figshare database (PacBio HiFi-reads from cod individuals is https://doi.org/10.6084/m9.figshare.24339823.v1).

## Author contributions

KSJ, RAA, WBR, OKT and JC conceptualized the study with input from CG. TJD. did DNA extractions with assistance from AK. RAA handled, processed and analyzed the data with input from WBR, OKT, JC, CG and KSJ. TJD analyzed the population data with contributions from RAA, HTB and MOSB. RAA wrote the manuscript with input from JC, KSJ, WBR, OKT, CG, SNKH, HTB, TJD, SJ, AK, and MOSB.

## Notes

https://doi.org/10.6084/m9.figshare.24339823.v1

### Competing Interest Statement

The authors have declared no competing interest.

