## Supplementary Figures for "Chromosomal inversions mediated by tandem insertions of transposable elements"

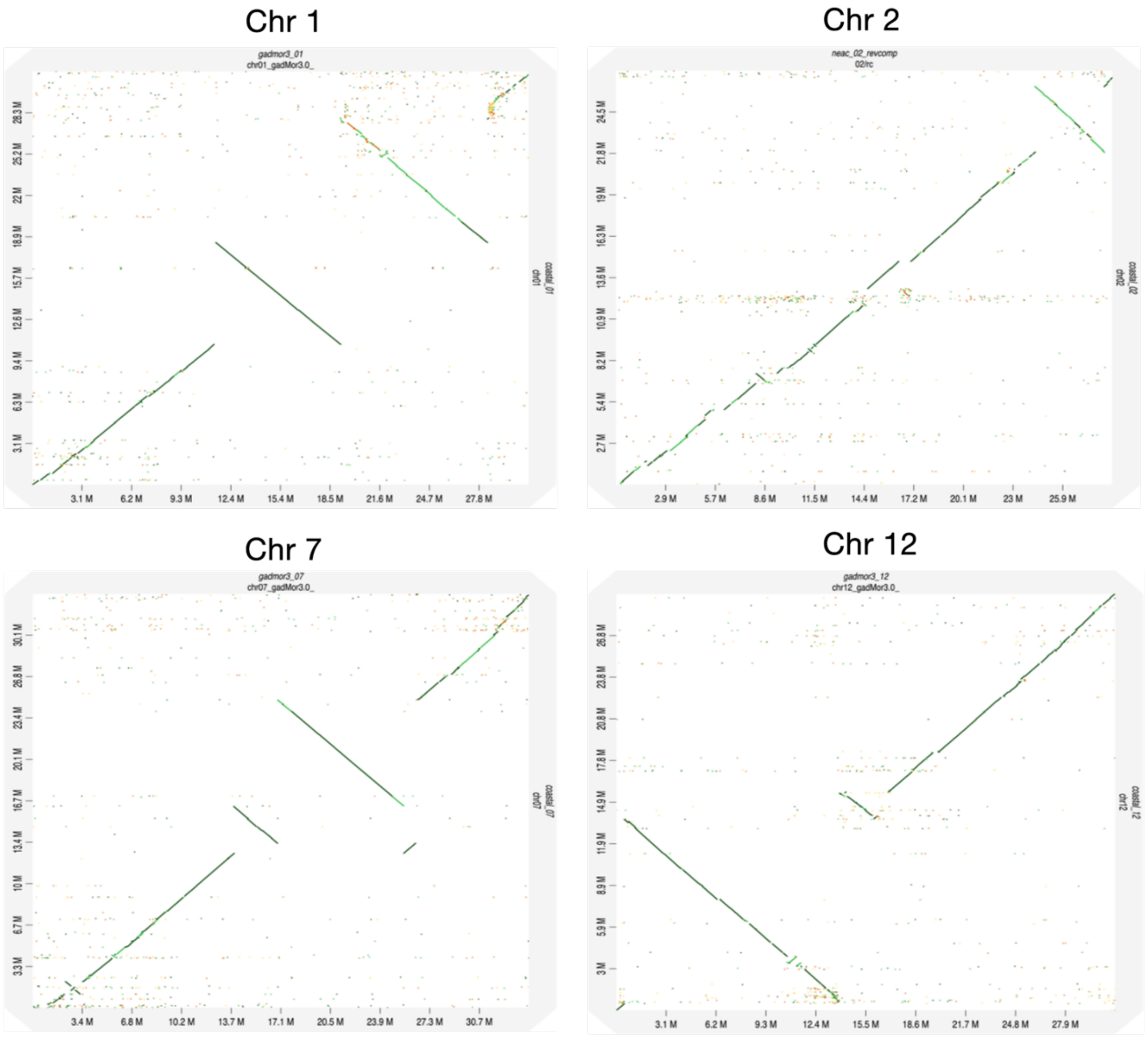


**Supplementary Figure S1: Dot-plots of chromosomes 1, 2, 7 and 12 in NEAC against NCC.** Dot-plot alignment of chromosomes 1, 2, 7 and 12 for NEAC (x-axis) against NCC (y-axis). Positions on the axes are shown in megabases (Mb).

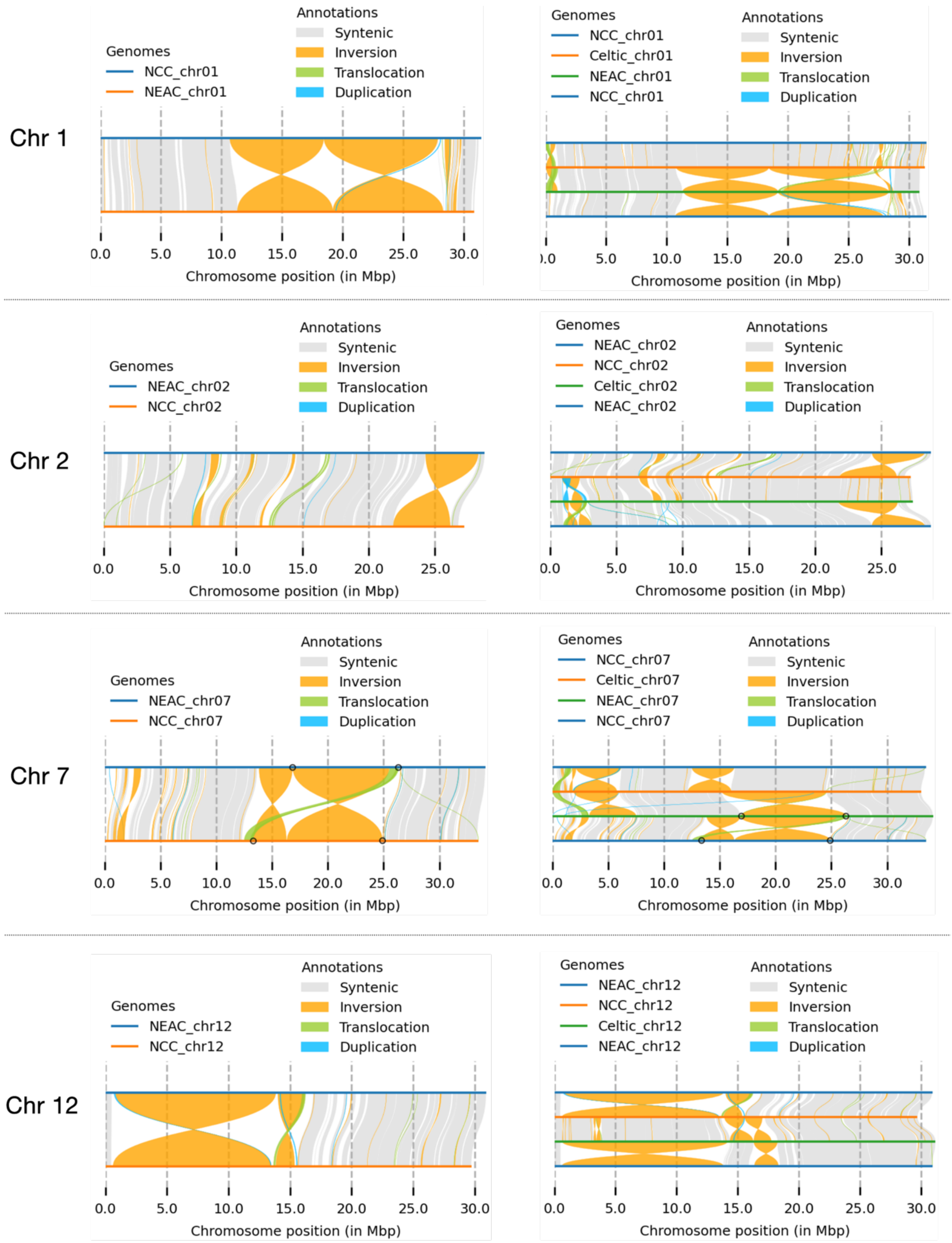


**Supplementary Figure S2: Synteny-plots of chromosomes 1, 2, 7 and 12 in NEAC against NCC and Celtic cod.** Synteny-plots are based on coordinates from mapping of scaffolds with minimap2 and variant-calling with SyRi. Reference genomes and structural variants are coloured according to legend. Positions on the x-axis are shown in megabases (Mb). For chromosome 7, the breakpoints of the single inversion used throughout the analyses are marked with black circles. These coordinates were evaluated as the most likely (true) breakpoint candidates based on synteny between the three assemblies (representing the loci from where the inversion breakpoints evolved).

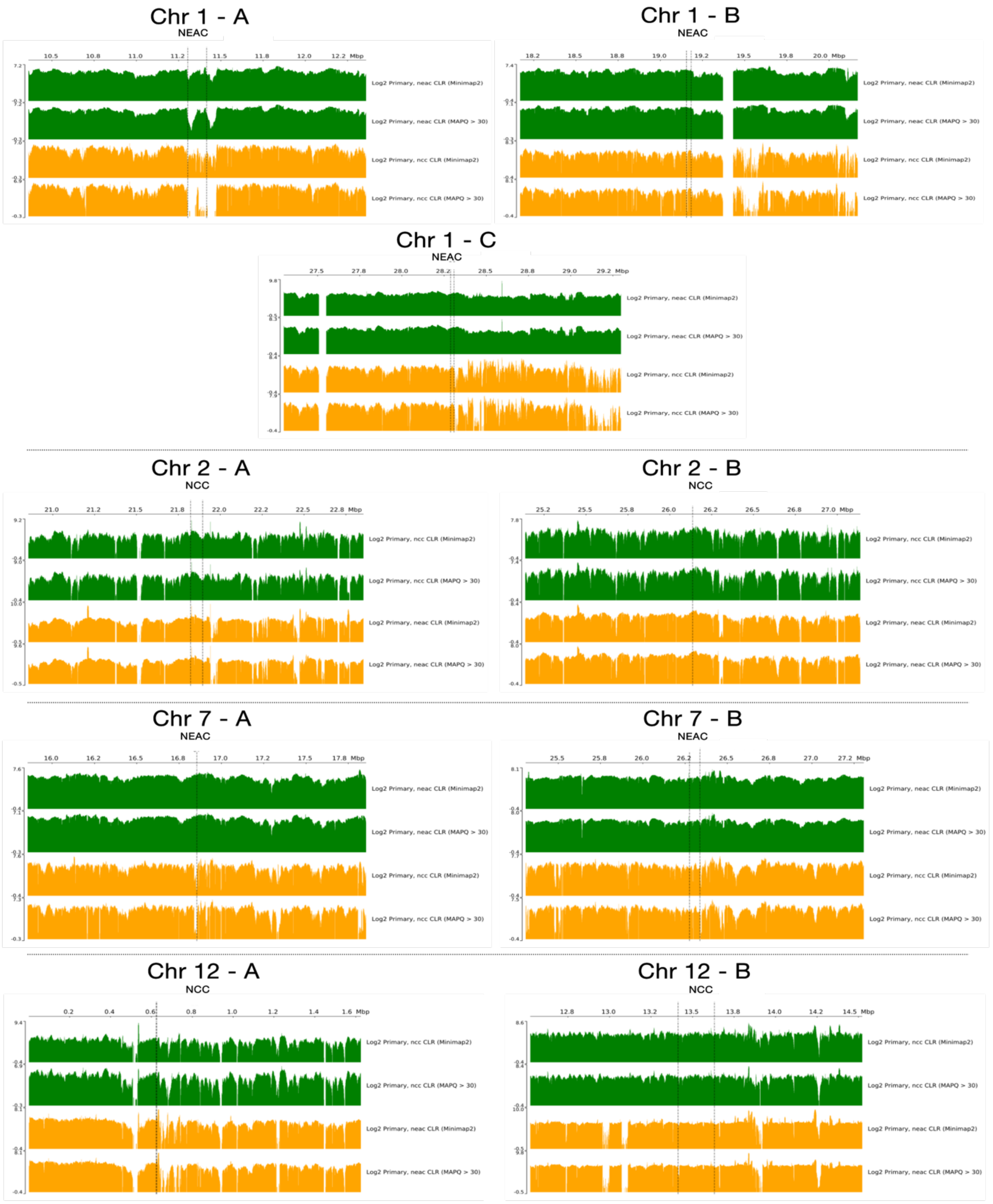


**Supplementary Figure S3: Read coverage plots of chromosomes 1, 2, 7 and 12 in NEAC and NCC.** Coverage plots showing PacBio CLR reads from NEAC and NCC assemblies mapped onto the reference genome carrying the derived (inverted) haplotype for each of the four chromosomes. Mapping of reads to the same assembly is shown in green (e.g., NEAC onto NEAC), while cross-mapping is in yellow (e.g., NCC onto NEAC). The y-axis displays Log2-coverage of primary reads, with tracks for mapping quality > 30 and without any quality cut-off. The x-axis shows positions in megabases (Mb) for regions around the breakpoints. Vertical dotted lines indicate initial breakpoint estimates from SyRi (see **Supplementary Figure S2**).

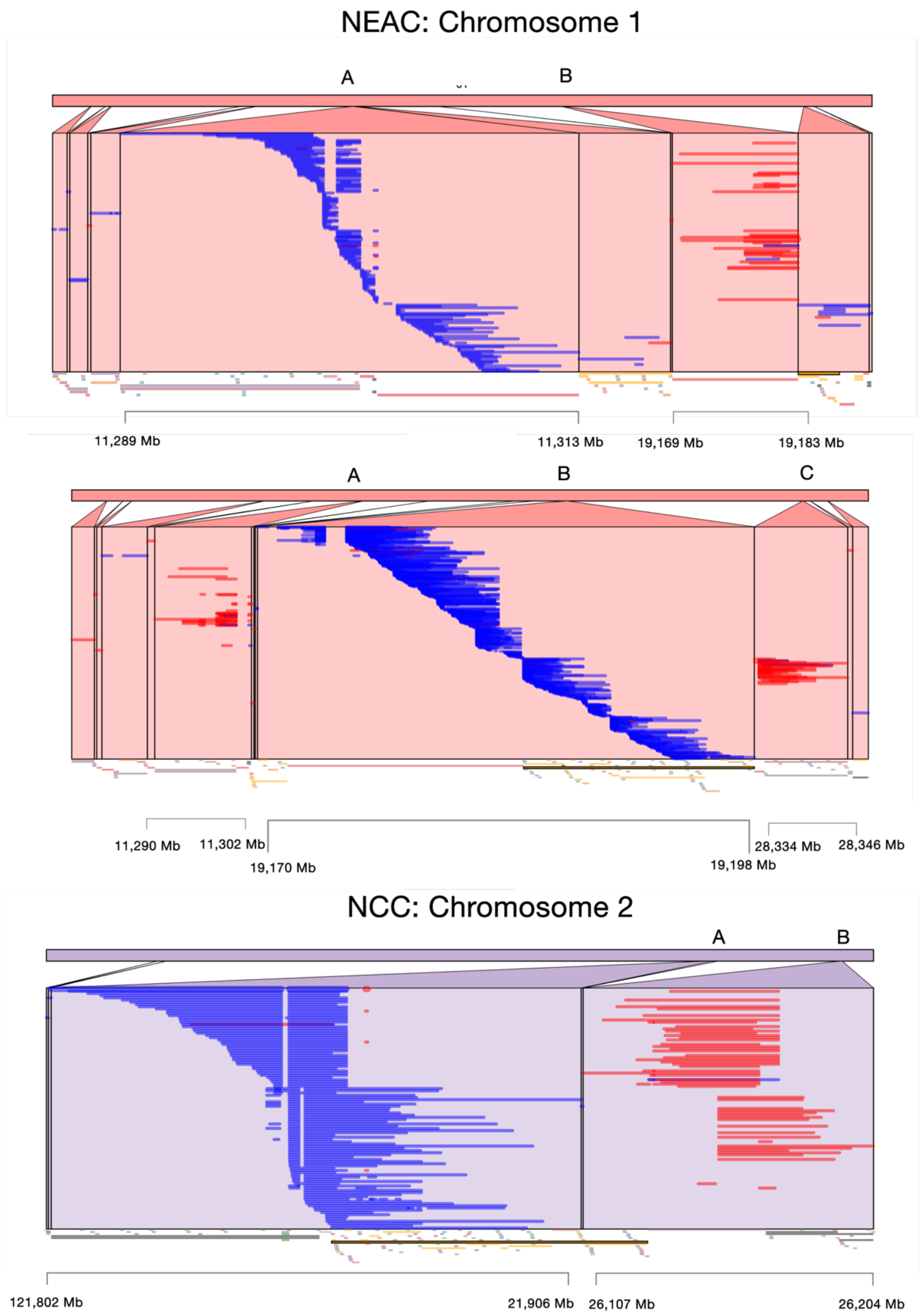


**Supplementary Figure S4: Mapping of raw reads on chromosomes 1, 2, 7 and 12 in NEAC and NCC.** Zoom-in on mapped (hard-clipped) reads from aligning long reads (NCC: CLR; NEAC: CLR, HiFi, CCS) to NEAC and NCC assemblies. The figures (NEAC: Chromosome 1 and NCC: Chromosome 2) only include cross-mapping of reads mapped onto the assemblies carrying the derived (inverted) haplotype for each of the four inversions. Boundaries of the breakpoint intervals were estimated using the edges of reads that mapped to two breakpoint locations in opposite directions. The tracks below the zoomed-in regions show structural variants as predicted by SyRi.

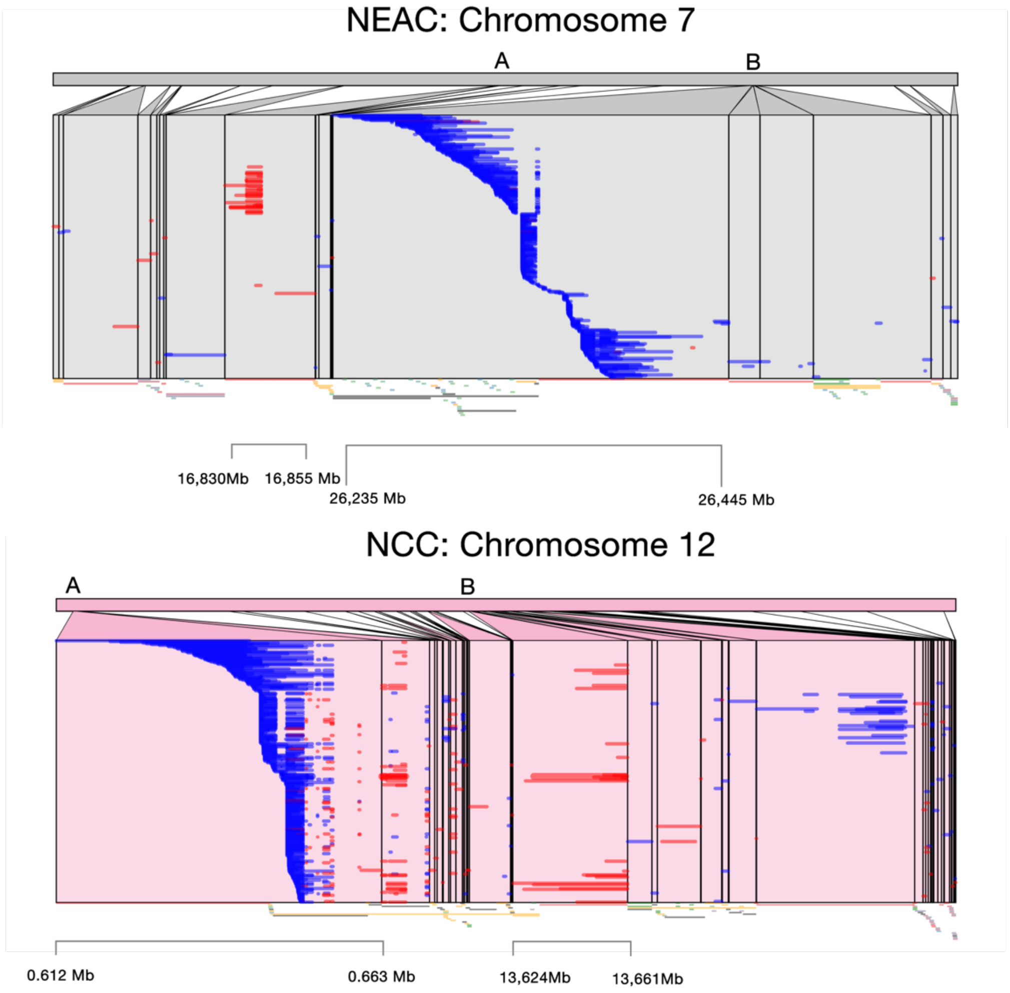


**Supplementary Figure S4: Mapping of raw reads on chromosomes 1, 2, 7 and 12 in NEAC and NCC (continued).** Zoom-in on mapped (hard-clipped) reads from aligning long reads (NCC: CLR; NEAC: CLR, HiFi, CCS) to NEAC and NCC assemblies. The figures (NEAC: Chromosome 7 and NCC: Chromosome 12) only include cross-mapping of reads mapped onto the assemblies carrying the derived (inverted) haplotype for each of the four inversions. Boundaries of the breakpoint intervals were estimated using the edges of reads that mapped to two breakpoint locations in opposite directions. The tracks below the zoomed-in regions show structural variants as predicted by SyRi.

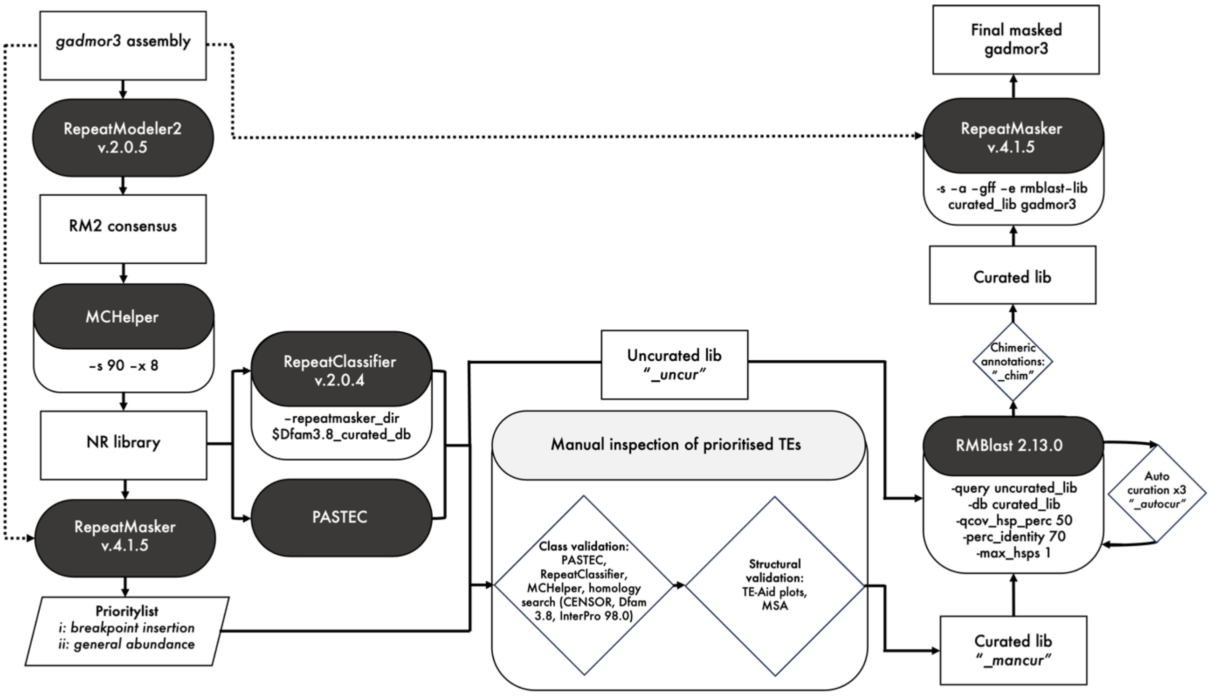


**Supplementary Figure S5: Workflow for *de novo* discovery, classification and annotation of TEs in Atlantic cod.** Step-by-step visualisation of our workflow for TE discovery, classification and annotation of the NEAC assembly (gadMor3.0). The programs used are shown in black rounded rectangles with settings below, input and output files are shown in white rectangles.

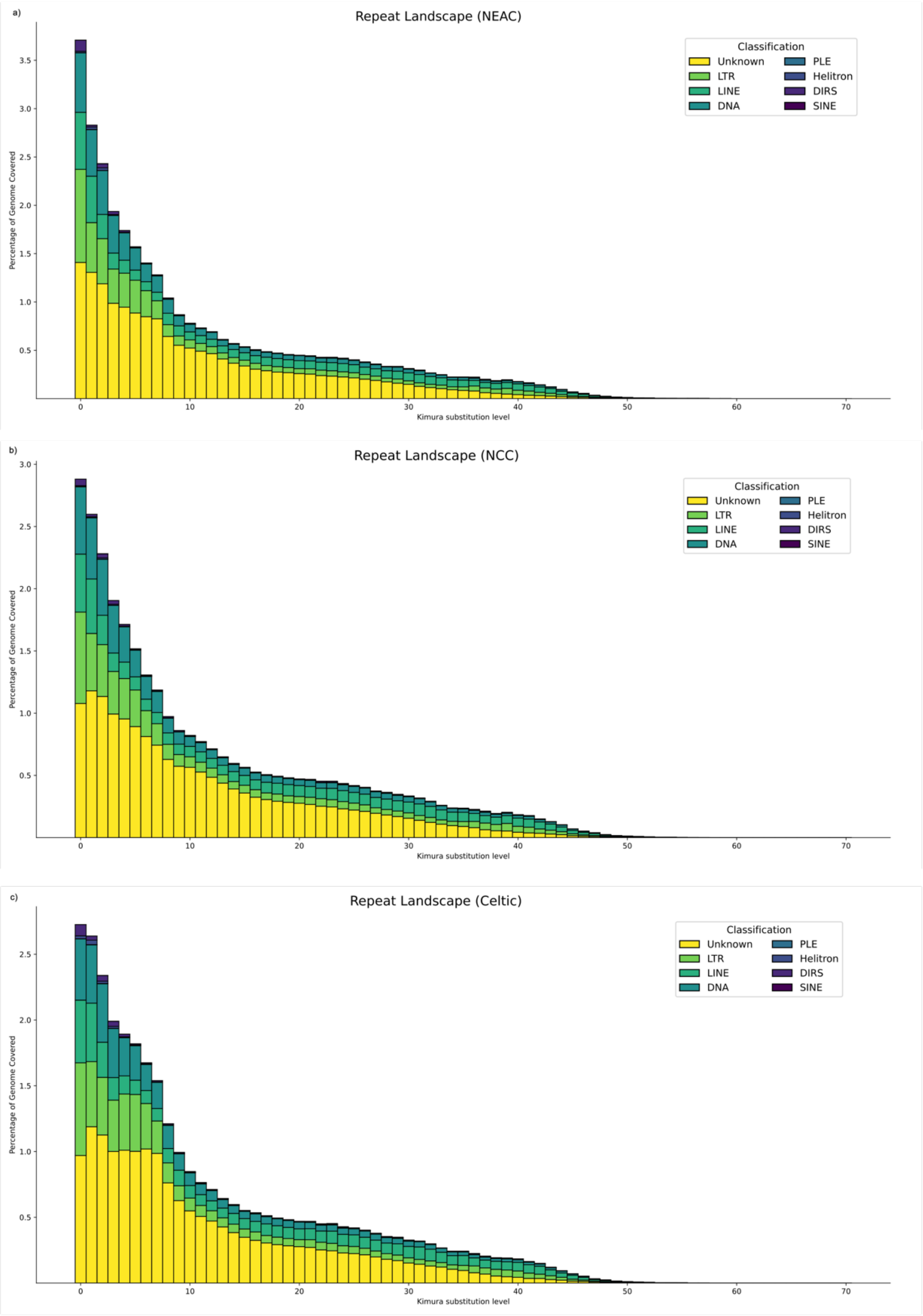


**Supplementary Figure S6: Repeat landscapes in Atlantic cod ecotypes.** The plots show percentage divergence of LTRs, LINEs, DNA elements, Penelope elements, Helitrons, DIRS, SINEs and unknown TEs from their consensus sequences on the x-axis (coloured according to legend) in **a,** NEAC, **b,** NCC, and **c,** Celtic cod. Genome coverage (%) is shown on the y-axis.

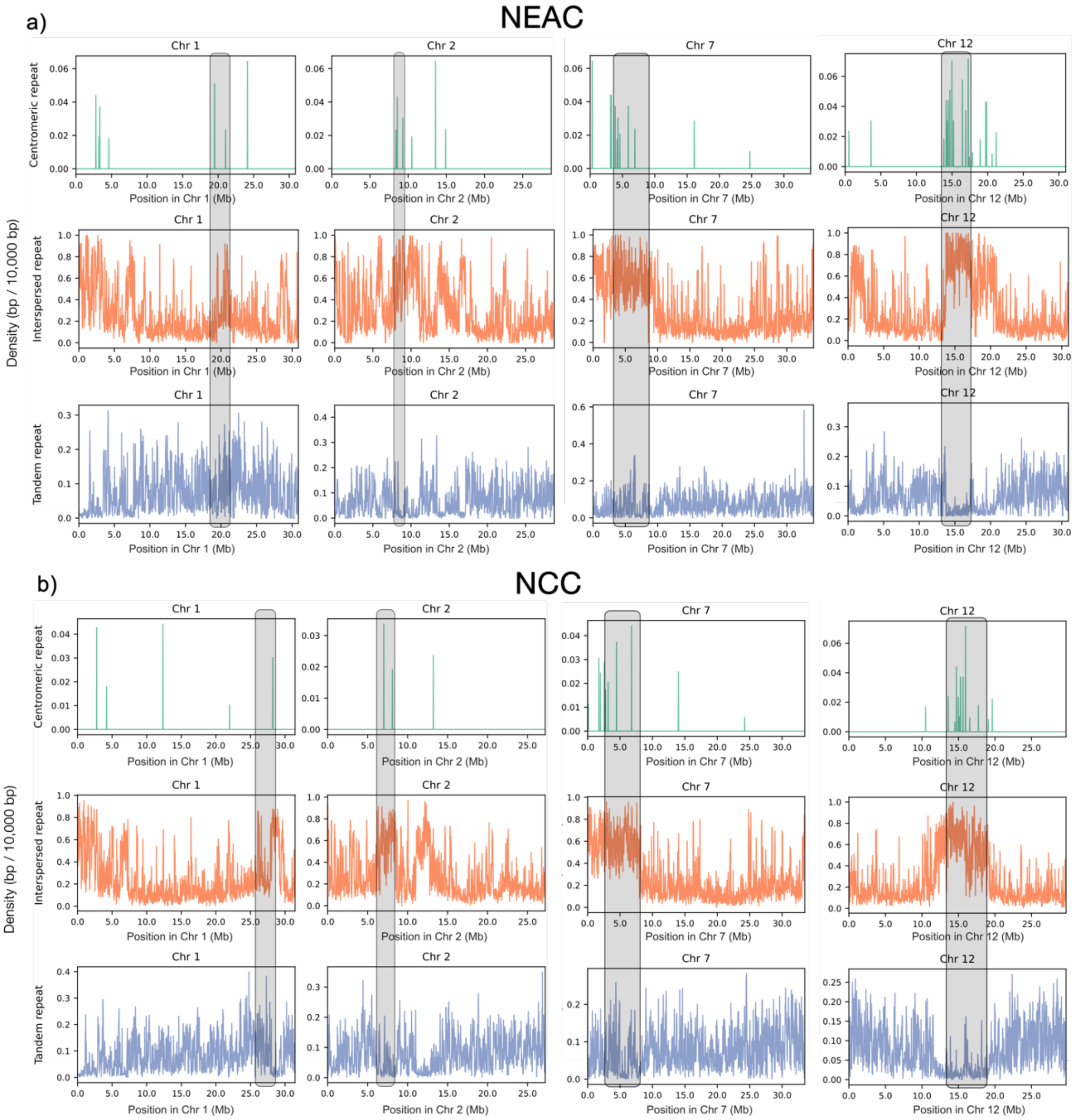


**Supplementary Figure S7: Putative centromeric repeats in Atlantic cod.** The top tracks (green) show mapping of the putative centromeric repeat from Kirubakaran et al. (2020) onto **a,** NEAC and **b,** NCC reference genomes. The middle tracks (orange) show the density of interspersed repeats within sliding windows of 10 kb in **a,** NEAC and **b,** NCC reference genomes. The bottom tracks (blue) show the density of putative tandem repeats within sliding windows of 10 kb in **a,** NEAC and **b,** NCC reference genomes. Putative centromeric regions are highlighted in grey for chromosomes 1, 2, 7 and 12. Positions on the x-axis are shown in megabases (Mb), density on the y-axis are shown as base pairs per 50,000 bp.

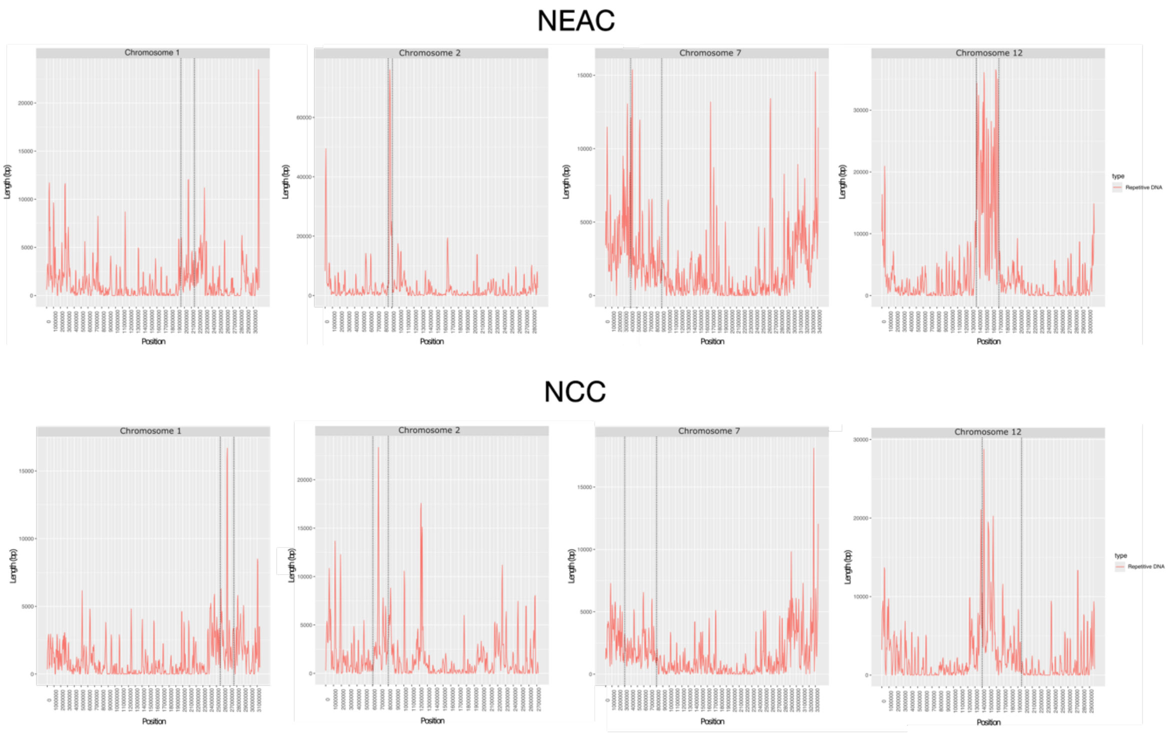


**Supplementary Figure S8: Putative centromere regions in chromosomes 1, 2, 7 and 12 NEAC and NCC.** Occurrence of best centromere repeat candidates in chromosomes 1, 2, 7 and 12, as predicted by CentroMiner. Chromosomal positions are shown on the x-axis (bp), length of repeats are shown on the y-axis. The vertical dotted lines indicate the boundaries of the putative centromere regions.

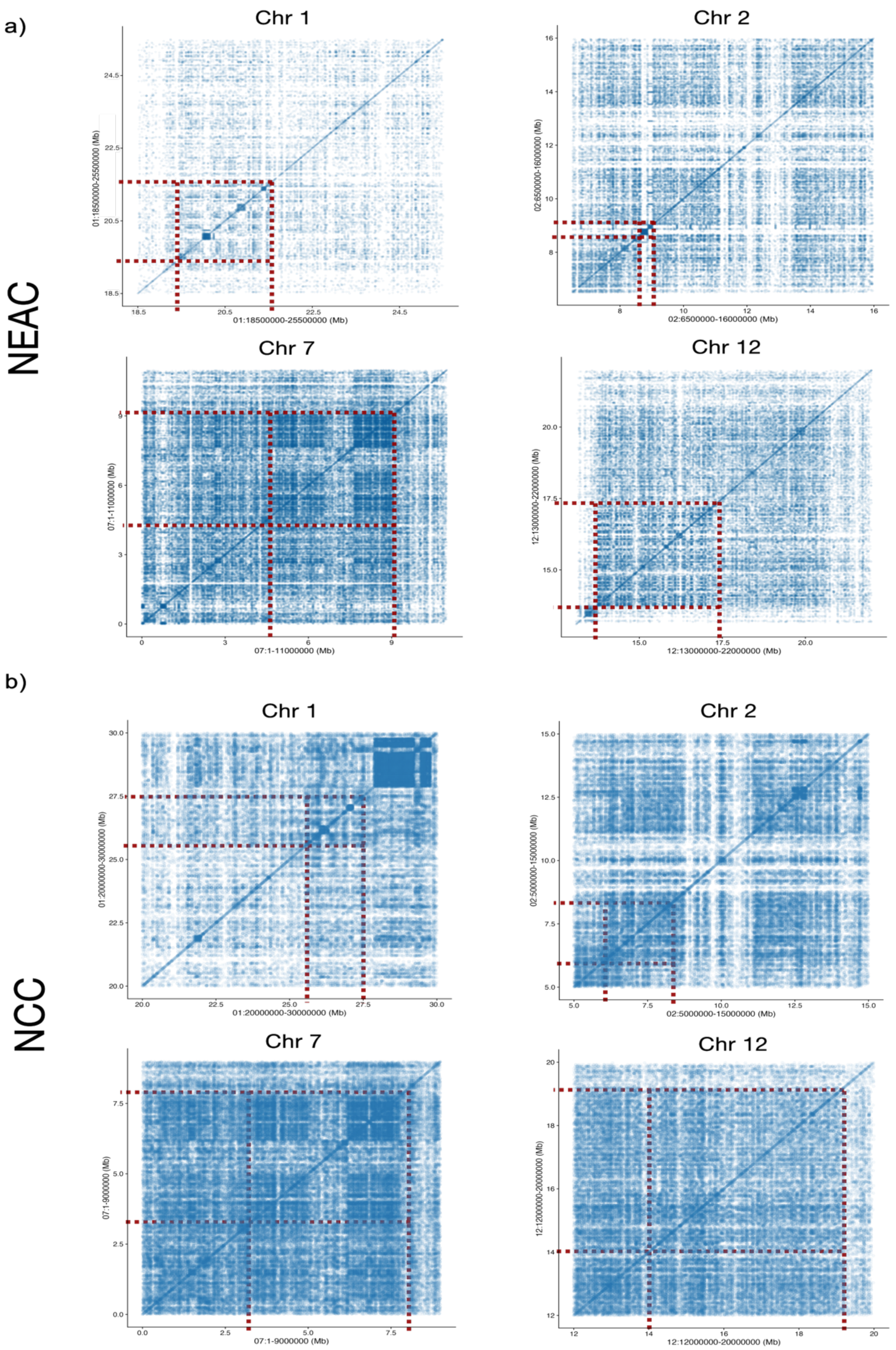


**Supplementary Figure S9: Putative centromere regions in chromosomes 1, 2, 7 and 12 NEAC and NCC.** Dot-plots of candidate centromere regions from **Supplementary Figures S7-S8**. The red dotted-lines indicate the boundaries of the putative centromeres used throughout the analyses. Positions on the axes are shown in megabases (Mb).

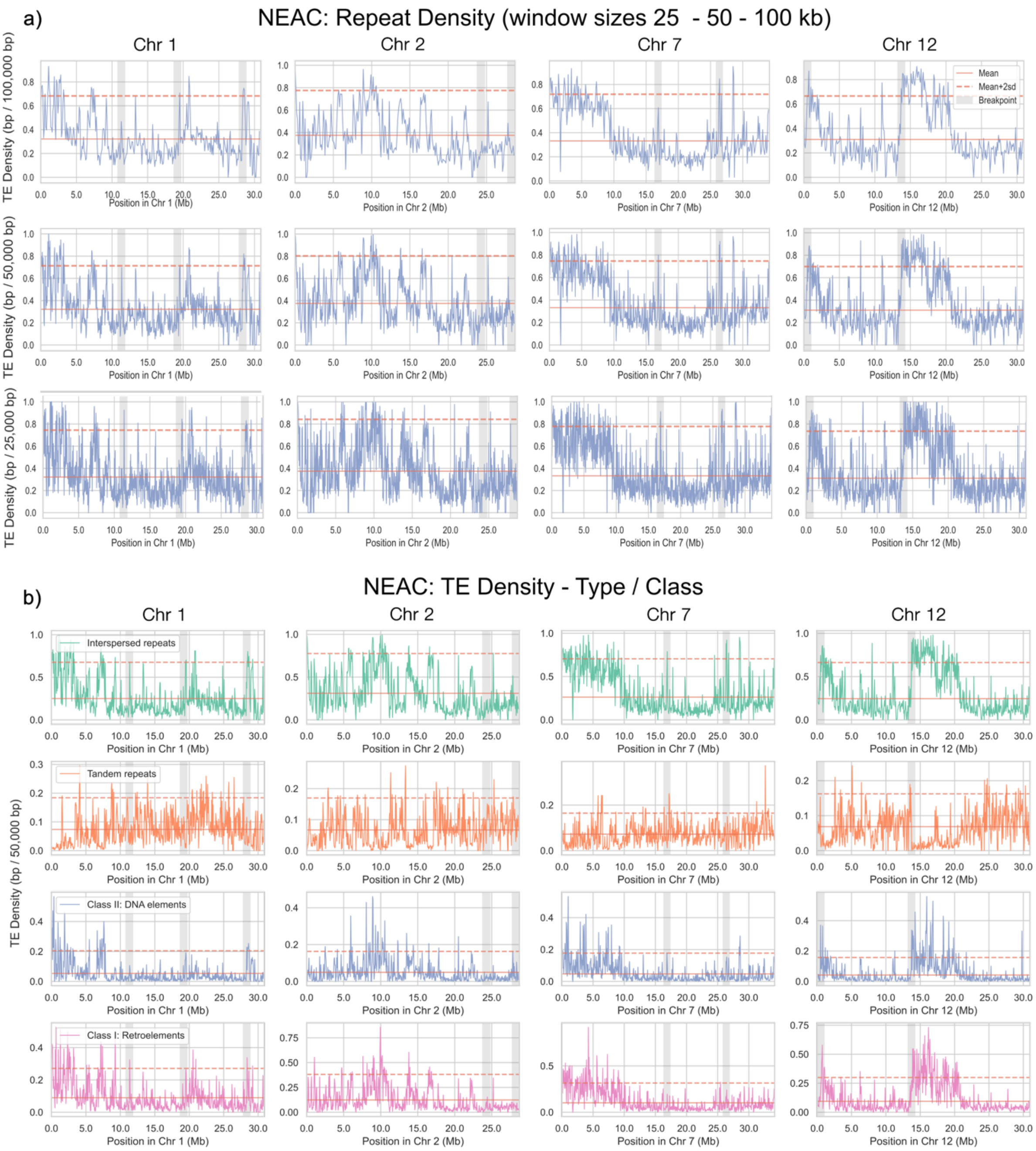


**Supplementary Figure S10: TE density on chromosomes 1, 2, 7 and 12 in NEAC and NCC.** **a,** TE density (blue lines) is calculated as TE coverage (bp) per non-overlapping sliding window of sizes 25 kb, 50 kb, and 100 kb along chromosomes 1, 2, 7 and 12 in NEAC. **b,** Repeat density (within 50 kb windows) across chromosomes 1, 2, 7 and 12 in NEAC, partitioned by interspersed repeats, tandem repeats, Class II: DNA elements and Class I: Retroelements. Red solid line: chromosome-specific mean of TE density. Red dashed line: two standard deviations above the mean density. Grey shaded areas: regions spanning the breakpoint (± 500 kb for visualisation). Chromosomes 1 and 7 are derived haplotypes in NEAC.

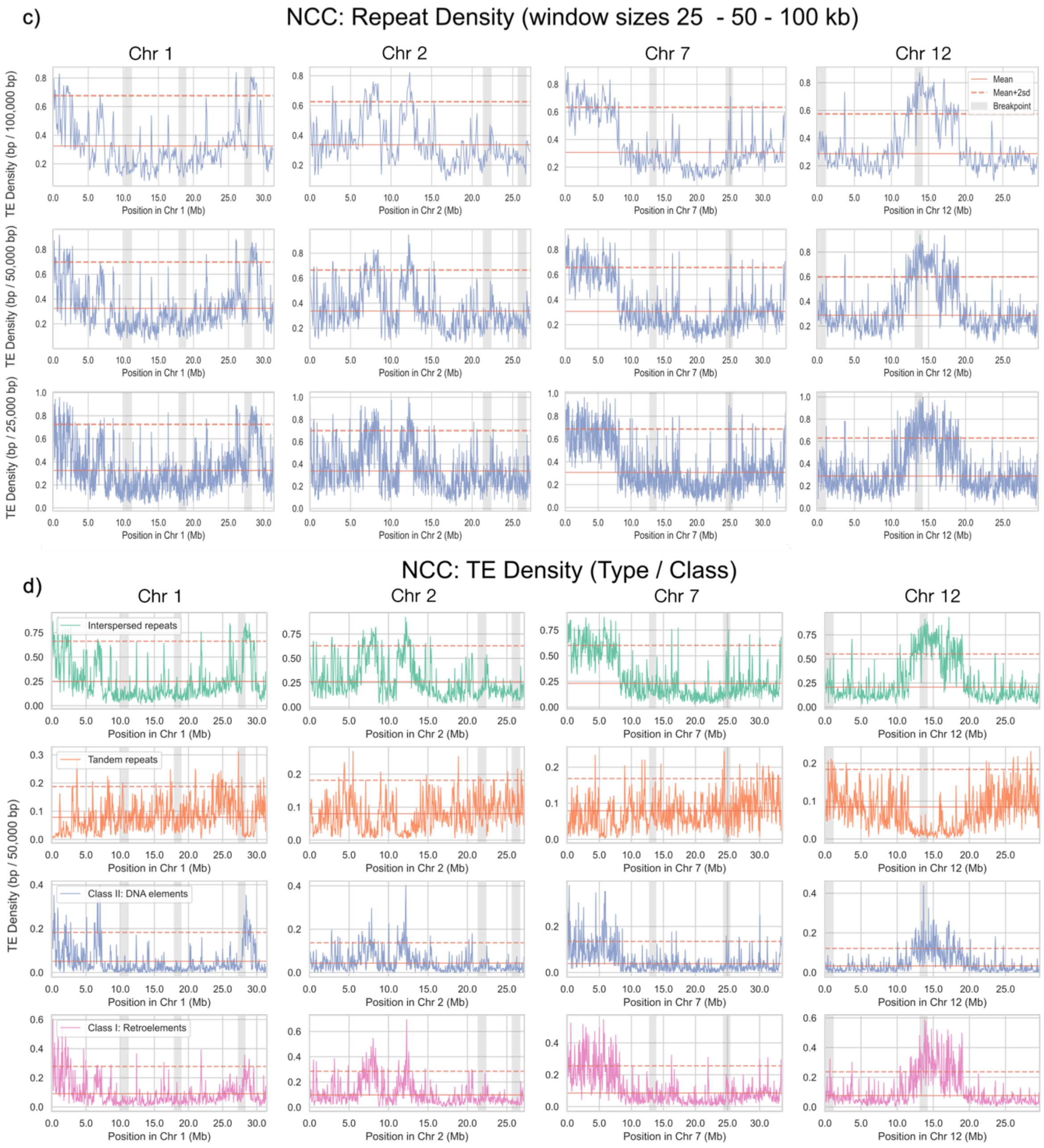


**Supplementary Figure S10: Putative centromere regions in chromosomes 1, 2, 7 and 12 NEAC and NCC (continued). c,** TE density (blue lines) is calculated as TE coverage (bp) per non-overlapping sliding window of sizes 25 kb, 50 kb, and 100 kb along chromosomes 1, 2, 7 and 12 in NCC. **d,** Repeat density (within 50 kb windows) across chromosomes 1, 2, 7 and 12 in NCC, partitioned by interspersed repeats, tandem repeats, Class II: DNA elements and Class I: Retroelements. Red solid line: chromosome-specific mean of TE density. Red dashed line: two standard deviations above the mean density. Grey shaded areas: regions spanning the breakpoint (± 500 kb for visualisation). Chromosomes 2 and 12 are derived haplotypes in NEAC.

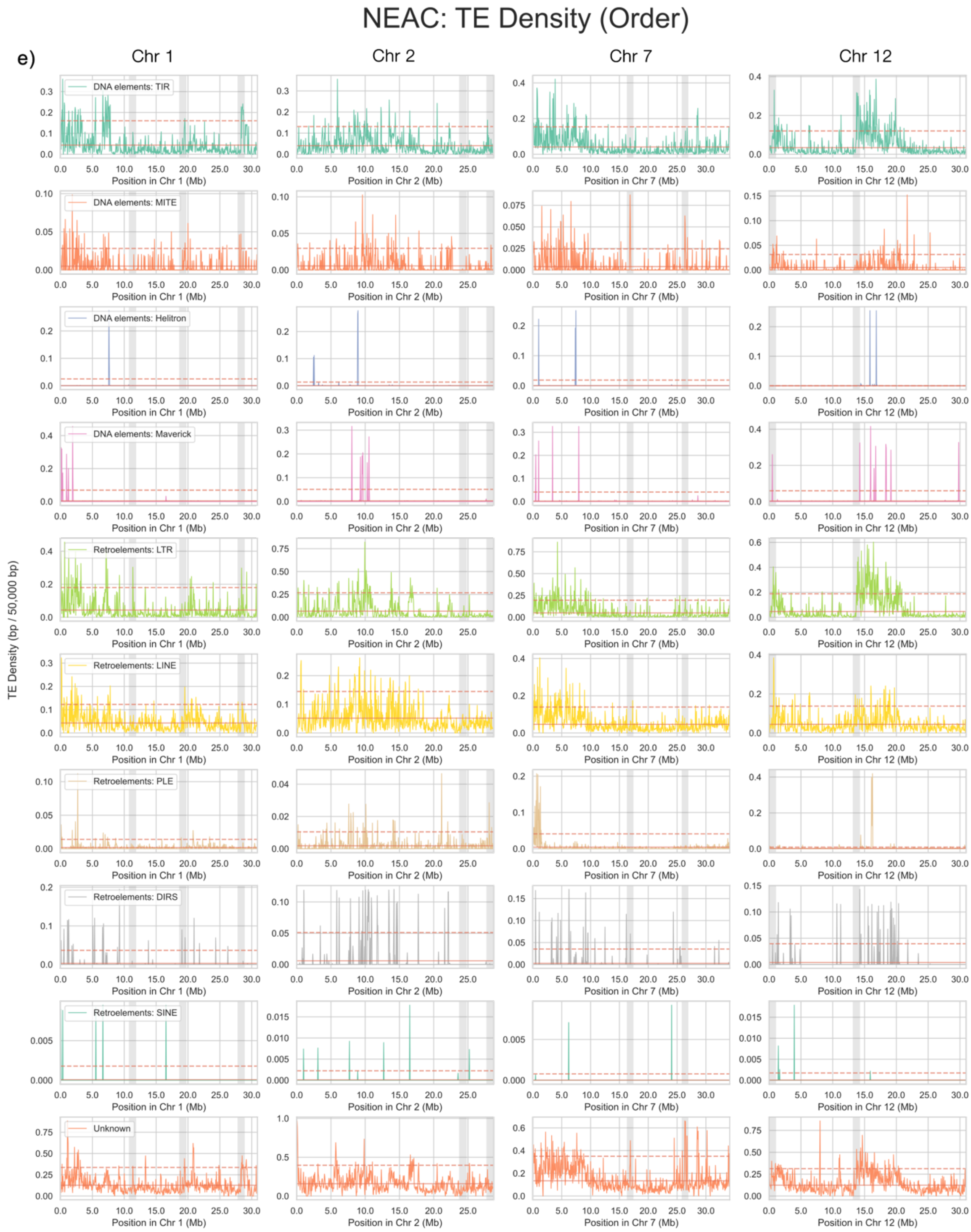


**Supplementary Figure S10: Putative centromere regions in chromosomes 1, 2, 7 and 12 NEAC and NCC (continued). e,** TE density (within 50 kb windows) along chromosomes 1, 2, 7 and 12 in NEAC, partitioned by TE orders (label and colour according to legend). Red solid line: chromosome-specific mean of TE density (excluding putative centromeres as estimated in **Supplementary Figures S7-S9**). Red dashed line: two standard deviations above the mean density. Grey shaded areas: regions spanning the breakpoint (± 500 kb for visualisation). Chromosomes 1 and 7 are derived haplotypes in NEAC.

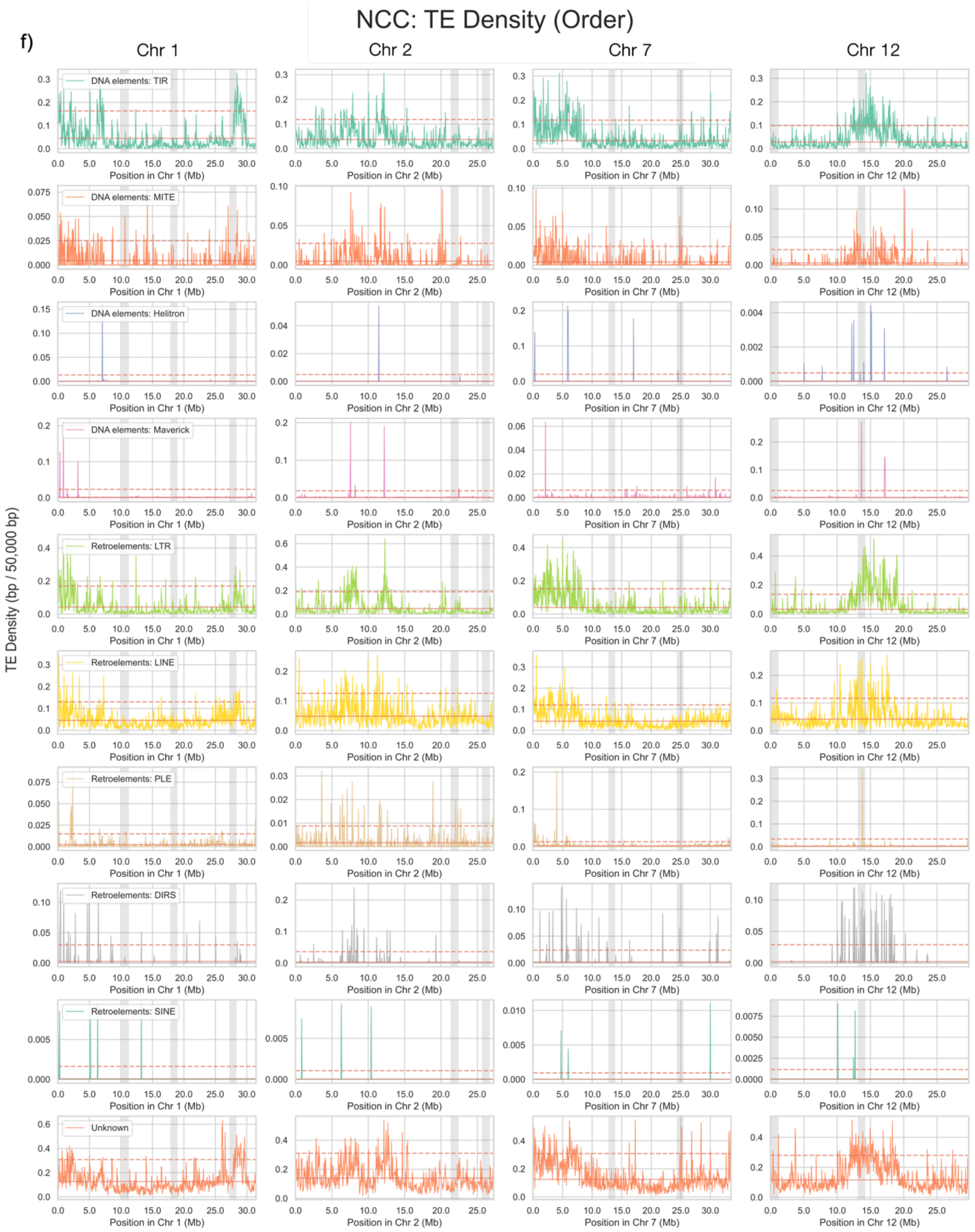


**Supplementary Figure S10: Putative centromere regions in chromosomes 1, 2, 7 and 12 NEAC and NCC (continued). f,** TE density (within 50 kb windows) along chromosomes 1, 2, 7 and 12 in NCC, partitioned by TE orders (label and colour according to legend). Red solid line: chromosome-specific mean of TE density (excluding putative centromeres as estimated in **Supplementary Figures S7-S9**). Red dashed line: two standard deviations above the mean density. Grey shaded areas: regions spanning the breakpoint (± 500 kb for visualisation). Chromosomes 2 and 12 are derived haplotypes in NCC.

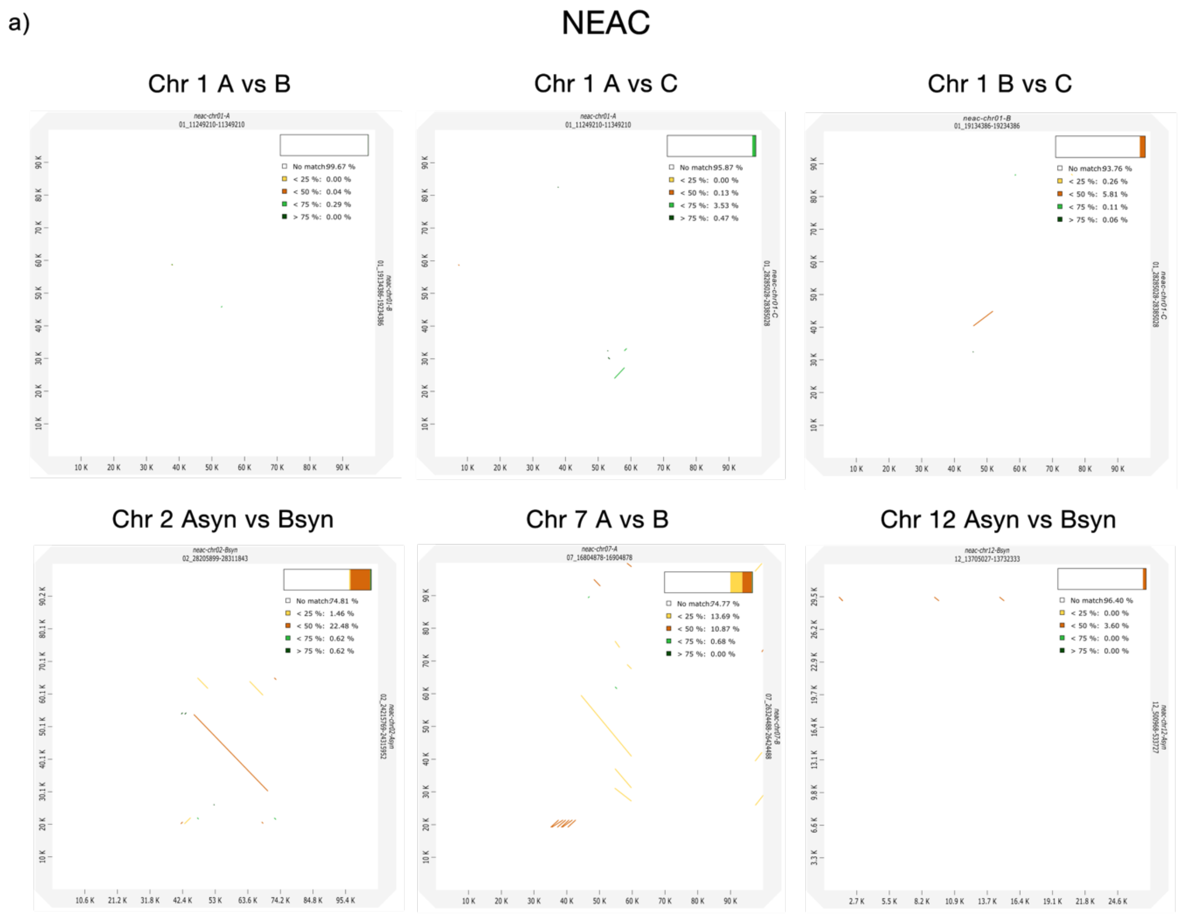


**Supplementary Figure S11: Dot-plot of breakpoint pairs in inversion haplotypes of NEAC and NCC.** Dot-plot alignment of all breakpoint pairs (± 50 kb) (chromosomes 1 and 7) and syntenic "breakpoint" pairs (chromosomes 2 and 12) in NEAC. Percentage identity of alignments are coloured according to legend. Positions on the axes are shown in kilobases (kb). Coordinates of the plotted regions are shown in the axis labels.

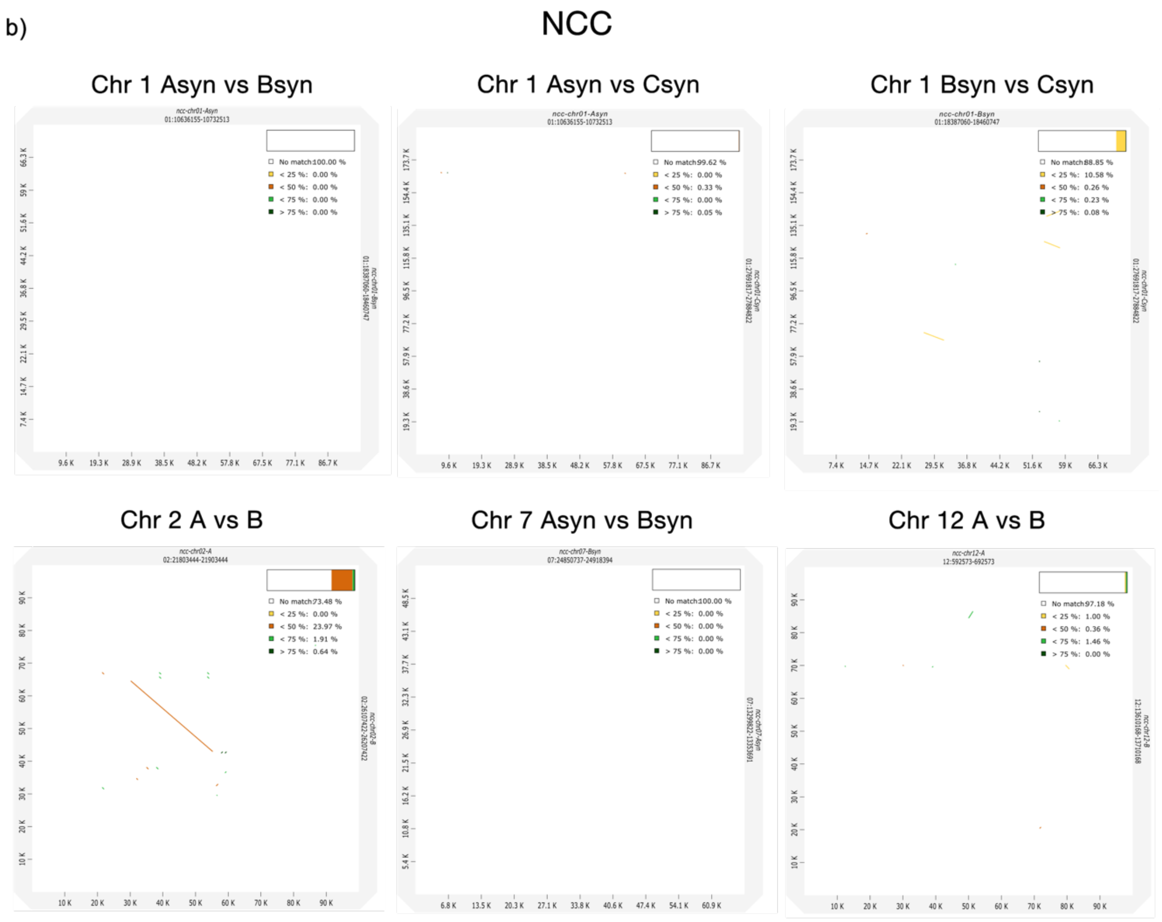


**Supplementary Figure S11: Dot-plot of breakpoint pairs in inversion haplotypes of NEAC and NCC (continued).** Dot-plot alignment of all breakpoint pairs (± 50 kb) (chromosomes 2 and 12) and syntenic "breakpoint" pairs (chromosomes 1 and 7) in NCC. Percentage identity of alignments are coloured according to legend. Positions on the axes are shown in kilobases (kb). Coordinates of the plotted regions are shown in the axis labels.

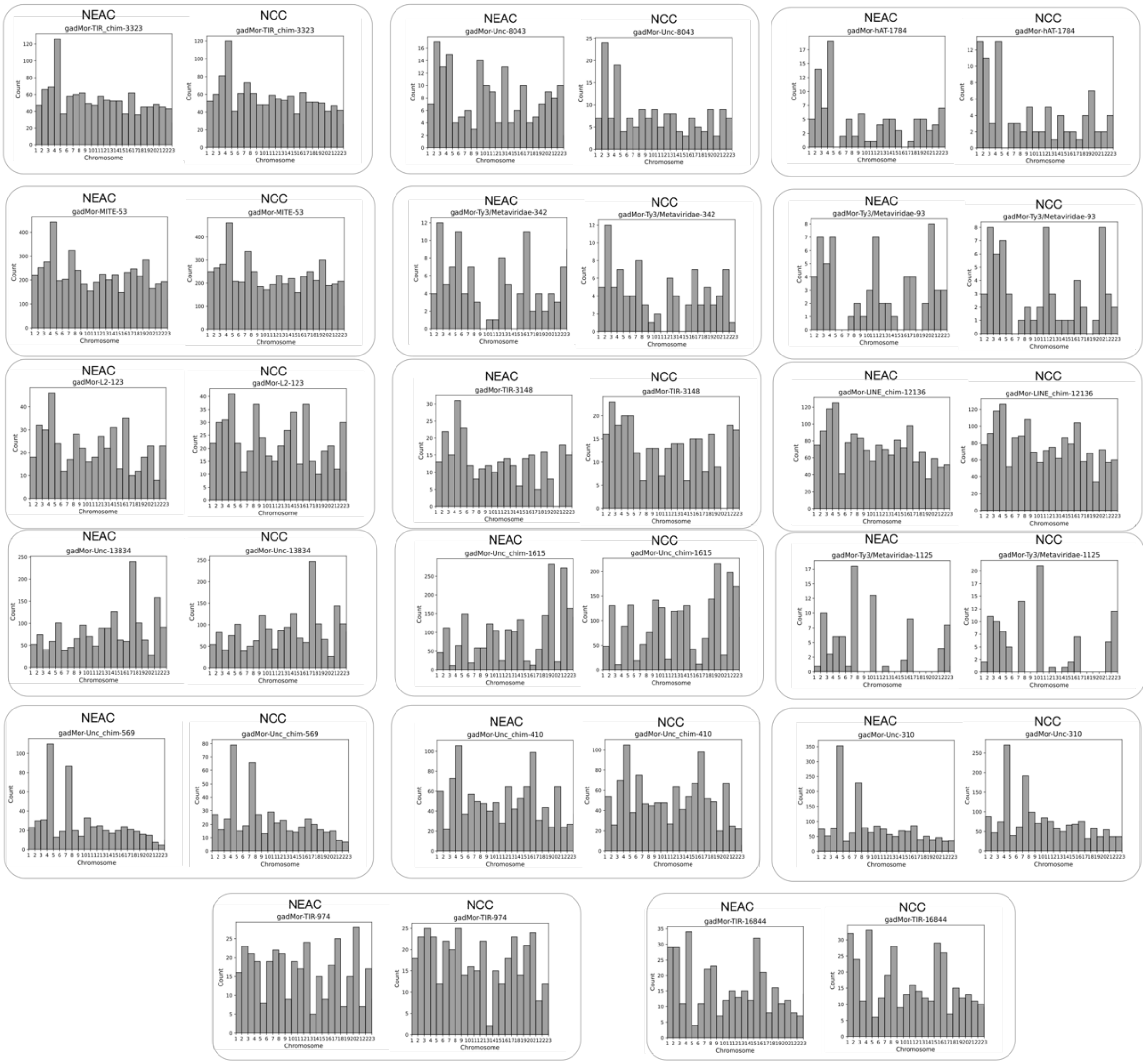


**Supplementary Figure S12: Genomic distribution of curated breakpoint TE families in NEAC and NCC.** The genomic distribution (insertions per chromosome) for each TE family that appears in breakpoint pairs.

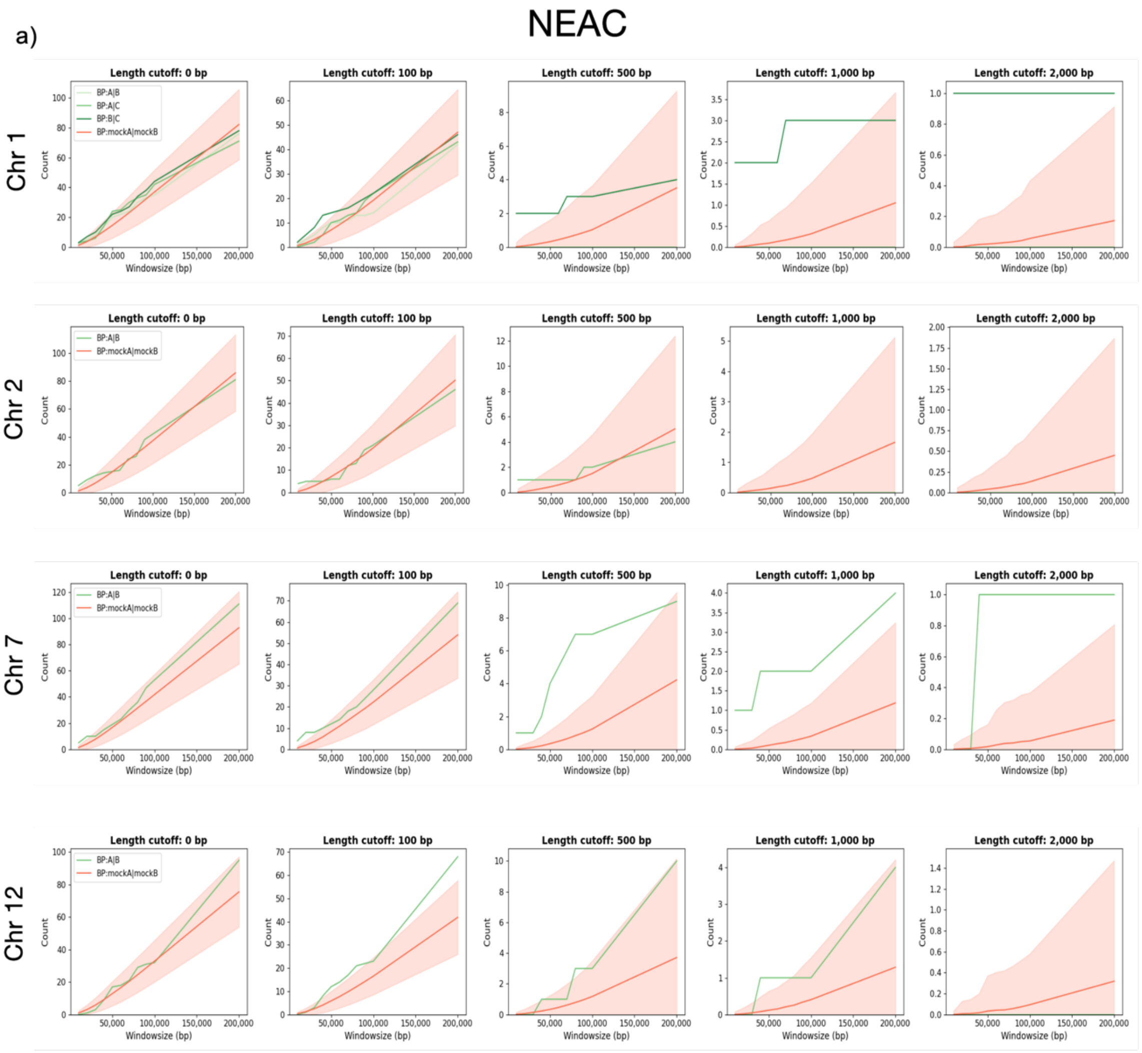


**Supplementary Figure S13: Probability of finding related TEs in breakpoints vs. random regions in chromosomes 1, 2, 7 and 12. a,** Plots for estimating the chance of finding related TEs in breakpoints compared to random region pairs in chromosomes 1, 2, 7 and 12 in NEAC. Count of related TEs (y-axis) with different max lengths (length cutoff: 0 bp, 100 bp, 500 bp, 1,000 bp and 2,000 bp) are plotted within window sizes of 10-100 kb (x-axis). Green lines show the observed count of related TEs within increasing window sizes around the true breakpoint pairs. Red lines show the average count of related TEs within 1,000 randomly distributed 'mock' breakpoint pairs. Pink shaded areas are standard deviations from the average TE count.

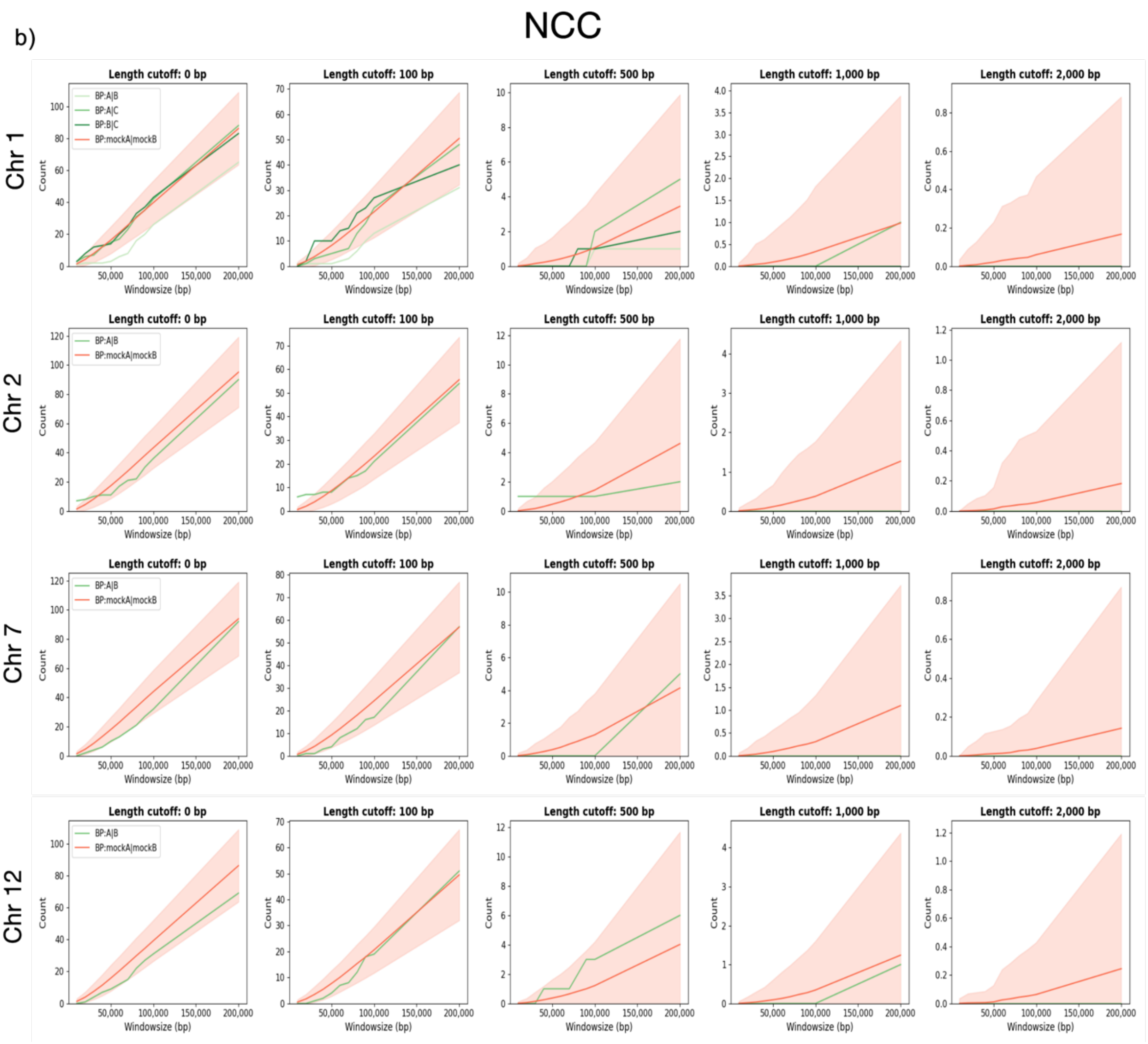


**Supplementary Figure S13: Probability of finding related TEs in breakpoints vs. random regions in chromosomes 1, 2, 7 and 12 (continued). b,** Plots for estimating the chance of finding related TEs in breakpoints compared to random region pairs in chromosomes 1, 2, 7 and 12 in NCC. Count of related TEs (y-axis) with different max lengths (length cutoff: 0 bp, 100 bp, 500 bp, 1,000 bp and 2,000 bp) are plotted within window sizes of 10-100 kb (x-axis). Green lines show the observed count of related TEs within increasing window sizes around the true breakpoint pairs. Red lines show the average count of related TEs within 1,000 randomly distributed 'mock' breakpoint pairs. Pink shaded areas are standard deviations from the average TE count.

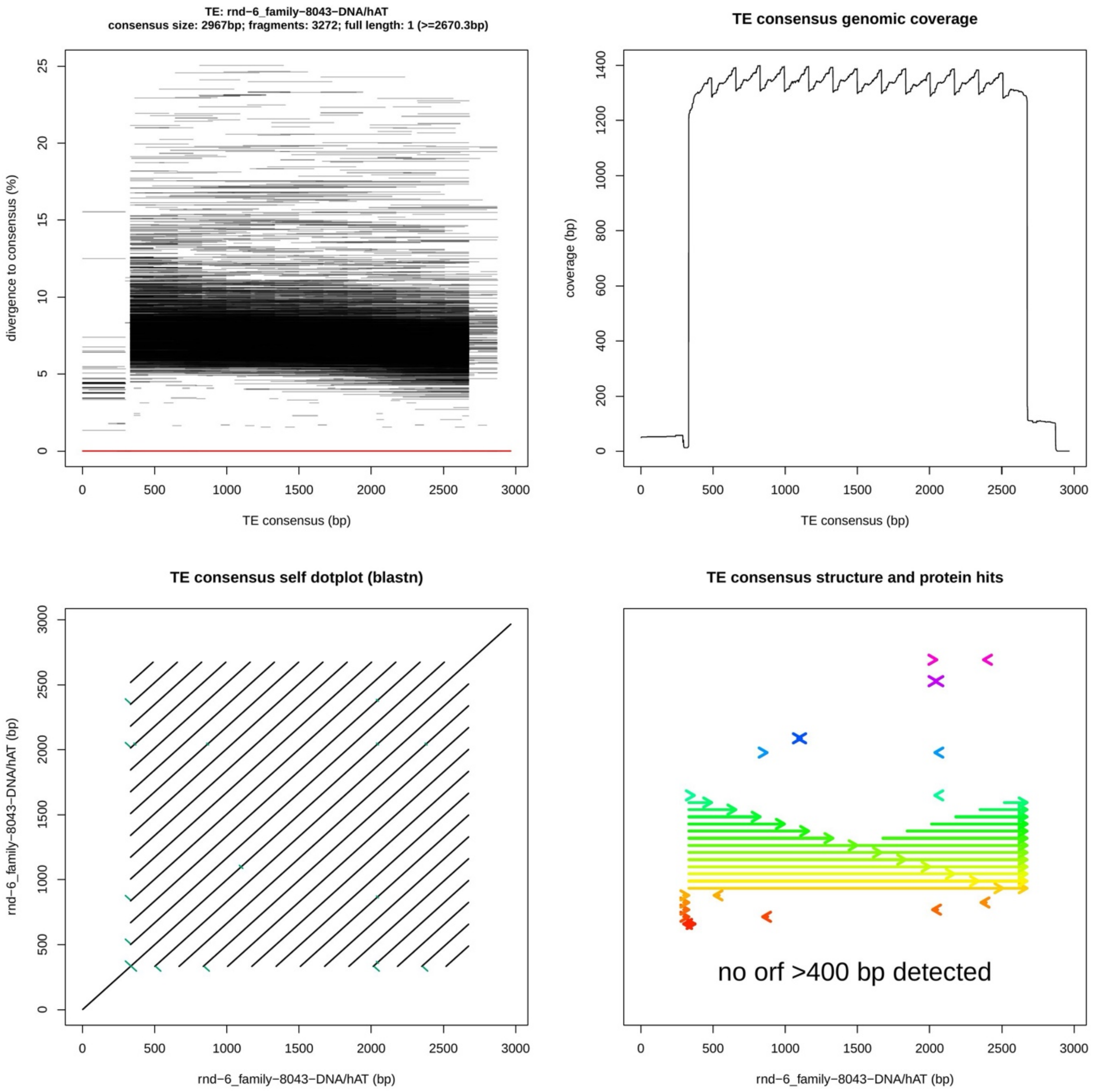


**Supplementary Figure S14: Structural analysis of gadMor-*hAT*-8043 with TE-Aid.** Output from TE-Aid (Goubert et al., 2022), visualising the gadMor-*hAT*-8043 family residing in the chromosome 1 and 7 breakpoints. **a,** Fragment and divergence plot after blasting consensus sequence against NEAC reference genome. Horizontal lines are genomic hits relative to the consensus sequence. Position on the y-axis represents divergence from the consensus. Red lines are full-length hits (spanning > 90% of the consensus). **b,** Sequence coverage of genomic hits from BLAST relative to the position along the TE consensus. **c,** Self alignment dot-plot of the TE consensus. **d,** Putative ORFs and corresponding peptides located in the TE consensus. Arrows represent micro-homologies and repetitive DNA from the self-alignment dot-plot in **(c)**.

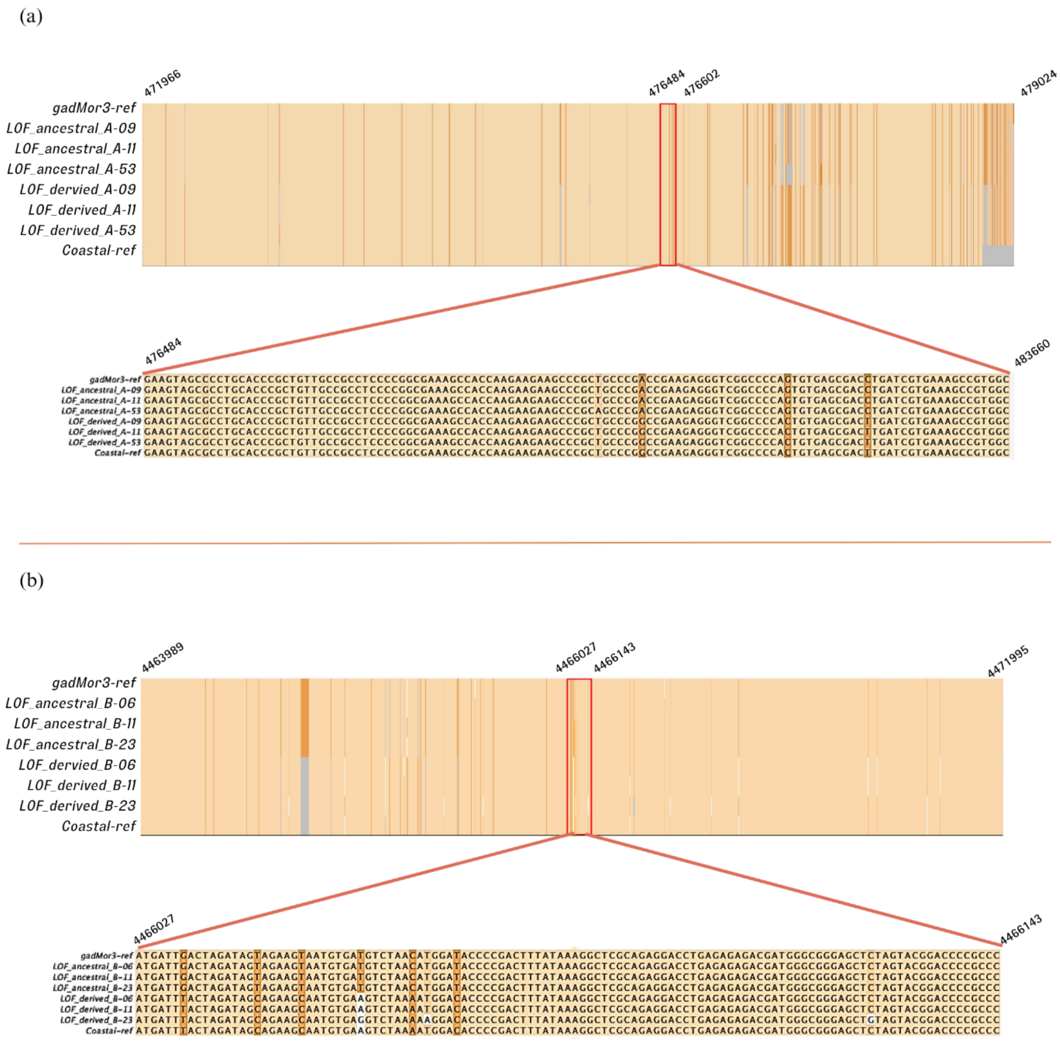


**Supplementary Figure S15: Breakpoint alignment of heterozygous individuals for the chromosome 2 inversion. a,** The alignment shows HiFi-sequences of chromosome 2 breakpoint A from heterozygous individuals with non-inverted (Asyn) and inverted (A) arrangements, along with the corresponding regions in the NEAC (top: NEAC-ref) and NCC (bottom: NCC-ref) reference genomes. **b,** Similarly, the alignment displays HiFi-sequences of chromosome 2 breakpoint B for heterozygous individuals. For both alignments, light beige indicates 100% identity among all eight sequences. Orange highlights sequence variations between the inverted and non-inverted sites. White represents variations between four or fewer sequences, while grey illustrates deletions. Both **a** and **b** use the NEAC coordinates as a reference. The alignments provide an overview of the entire HiFi-sequenced regions, with a detailed alignment shown in the highlighted red box.

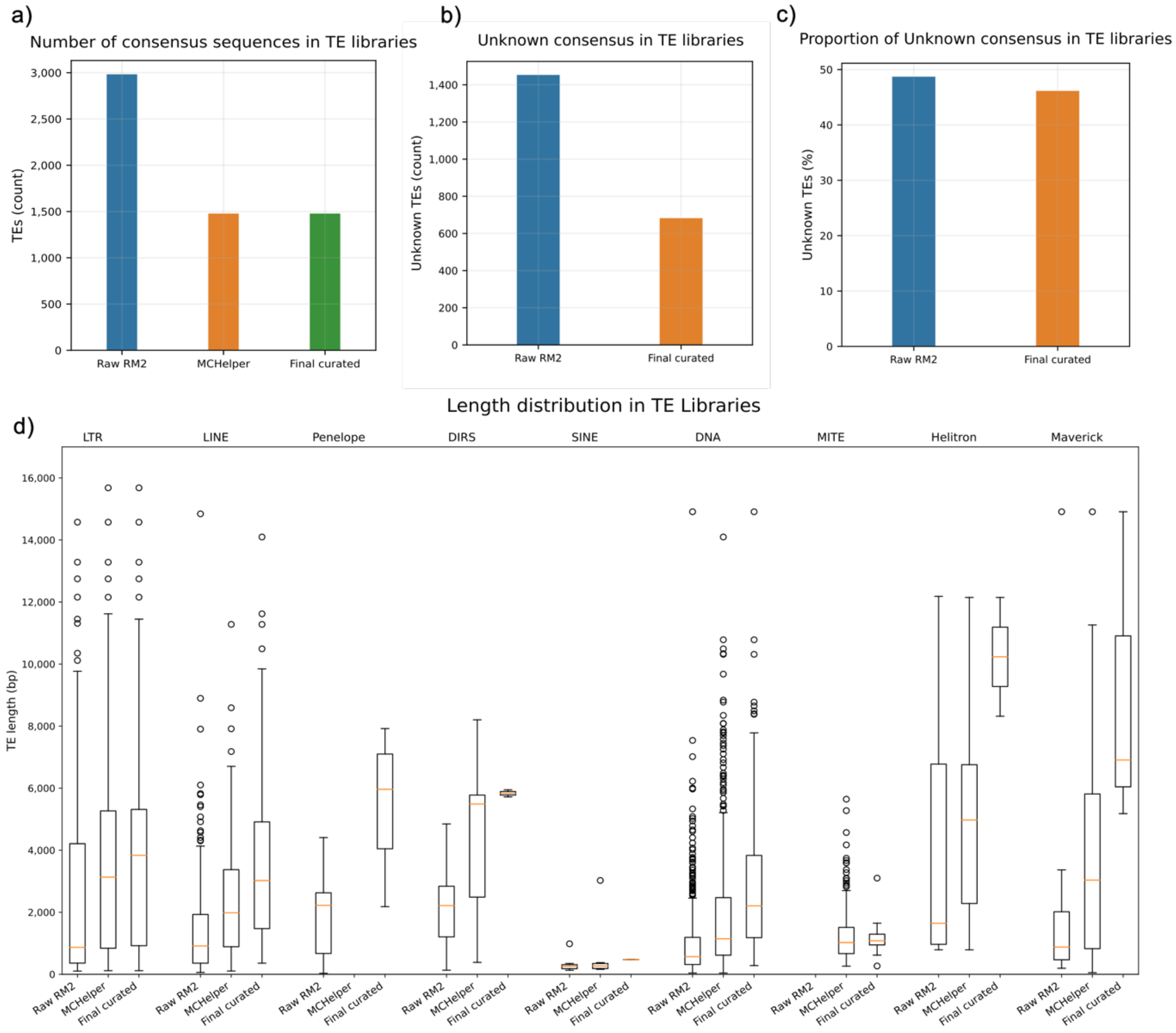


**Supplementary Figure S16: Comparison of raw, autocurated and manually curated TE library for *G. morhua*. a,** Total number of consensus sequences in the raw library (RM2), autocurated library from MCHelper and final curated TE library. **b,** Total number of unknown (unclassified) TEs in the raw (RM2) and final curated TE libraries. **c,** Proportion of unknown (unclassified) TEs in the raw (RM2) and final curated TE libraries. **d,** Boxplot of the length distribution of consensus sequences from different TE orders in the raw library (RM2), autocurated MCHelper library and final curated TE library. Length on the y-axis in shown in base pairs (bp).

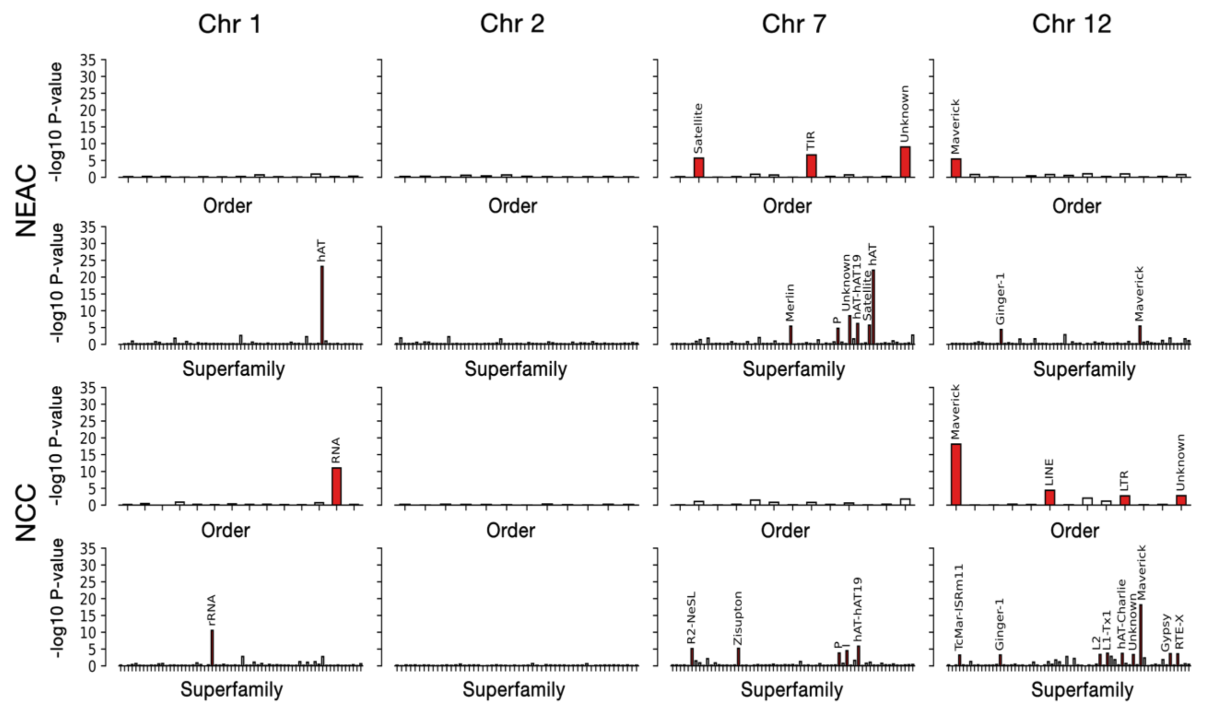


**Supplementary Figure S17: TE enrichment in breakpoints vs. non-breakpoint regions for uncurated TE library.** TE orders and superfamilies associated with breakpoint regions using a student's t-test. TE orders and superfamilies are shown on the x-axis, and -$\log_{10}$ p-values on the y-axis for chromosomes 1, 2, 7 and 12 in NEAC and NCC. Red bars indicate TE orders and superfamilies with significantly elevated densities within breakpoint regions.
